# *Tangerine:* a new family of *Starships* from lichen-forming fungi

**DOI:** 10.1101/2025.11.25.690456

**Authors:** Gulnara Tagirdzhanova, Noah E. Brown, Angus H. Bucknell, Ellen S. Cameron, Robert D. Finn, Mark Blaxter, Megan C. McDonald, Emile Gluck-Thaler, Nicholas J. Talbot

## Abstract

Lichens are symbiotic associations between filamentous fungi and photosynthetic micro-organisms, such as green algae and/or cyanobacteria, that result in a single anatomically-complex structure that can thrive in environments inhospitable to most organisms, including arctic tundra, high mountains, and deserts. Recent evidence suggests that lichens may be even more complex than previously appreciated, containing multiple microbial constituents, but how genomes of the principal fungal symbiont (which provides the majority of biomass in lichen tissue) have been shaped during evolution is largely unexplored. Recently, giant transposable elements called *Starships* have been found in many genomes of filamentous fungi, but to which extent they occur in lichen-forming fungi is not known. In this report, we describe a *Starship* element from the lichen fungus *Xanthoria parietina*. This element, named *Tangerine*, contains several genes that have signatures of horizontal gene transfer from non-lichen-forming fungi, most likely from black yeasts of the Chaetothyriales, that are often lichen-associated. Repetitive sequences carried by *Tangerine*, and found in other sites in *Xanthoria* genomes, are affected by repeat-induced point mutation (RIP), a mechanism of genome defense against transposable elements, consistent with fungal sexual reproduction which always precedes new lichen formation by *Xanthoria*. *Tangerine*’s “captain” belongs to a newly defined family of tyrosine recombinases specific to lichen-forming Lecanoromycetes. Several other captain clades have signatures of horizontal gene transfer between distantly related lichen-forming fungi and non-mycobiont lichen-associated fungi. We speculate that *Starships* may play a significant, yet hitherto unrecognized role, in lichen genome evolution and provide a roadmap for further investigation.

## Introduction

The lichen symbiosis is one of the most common, yet poorly understood fungal lifestyles. Lichens are an association between a hyphal fungus (the mycobiont) and a photosynthetic microorganism (the photobiont), in which the majority of the biomass comes from the fungal symbiont (1) but the overall shape and organisation of the lichen thallus is much more complex than either partner alone. Recent findings also suggest that lichen thalli can in addition contain many more taxa than the photobiont and the mycobiont, including additional fungi (sometimes referred to as endolichenic or lichenicolous fungi) and bacteria (2–4). Lichen-forming fungi account for approximately 17% of all described fungal species and 27% of ascomycetes (5). The majority of mycobionts are filamentous fungi belonging to the Pezizomycotina and chiefly to the class Lecanoromycetes, which consists of mostly lichen-forming fungi and is a sister clade to Eurotiomycetes (5, 6). Despite being ubiquitous in terrestrial ecosystems, lichens remain very poorly understood and much is unknown about the biology and evolution of these symbiotic associations. Lichen-forming fungi are challenging to isolate in axenic culture and notoriously slow-growing (7), which complicates lichen research – including the production of genomic data. The class Lecanoromycetes, for instance, has three times as many described species as the Eurotiomycetes, but the International Nucleotide Sequence Database Collaboration currently contains six times fewer Lecanoromycetes genomes, the vast majority of which are extracted from short-read metagenomic data and are therefore highly fragmented. Understanding genome evolution in lichen-forming fungi is therefore severely limited by available high quality genome data.

Transposable elements (TEs) are genetic units that may change their own position within their host’s genome (8). Found across all domains of life, TEs are thought to have a profound impact on their host’s biology by both mediating host evolution and inducing genomic instability (9–11). Recently, a novel group of TEs, known as the *Starships*, have been reported and implicated as vectors of fungal-fungal horizontal gene transfer (HGT) (12). Unlike other types of TEs, *Starships* are vast in size and carry variable sets of genes as cargo, many of which encode adaptive phenotypes (13, 14). Additionally, *Starships* carry a more conserved set of four “auxiliary” genes which are thought to be beneficial to the maintenance of the TE. The result of carrying both sets of genes is that *Starships* are significantly larger than other transposons, often surpassing 100 kb in size and containing between ten to several hundred genes (15). Very recent evidence has experimentally demonstrated *Starships’* transposition and their capacity to horizontally transfer between distantly related fungal species (16). While the mechanism of *Starship* transposition is unknown, it is driven by a tyrosine recombinase (tyrR) known as the ‘captain’. This tyrR is always located as the first gene at the 5’-end of the *Starship* element. *Starships* occur exclusively in Pezizomycotina fungi, the most species-rich group of fungi that includes over 90,000 species with ecologies ranging from saprotrophs to plant symbionts to animal and plant pathogens (17). *Starships* have been proposed to shape the evolution of Pezizomycotina fungi by constituting a large and mobile compartment of the genome and might in part be responsible for rapid and repeated evolution of ecological strategies observed in filamentous fungi (18).

Detection of *Starships* is challenging due to their large size and low copy number (15). Existing tools for *Starship* annotation, starfish (15) and stargraph (19), rely on aligning contiguous genome assemblies from the same species and identifying gaps in the alignment that contain tyrR-encoding genes. Because only a handful of high-quality genomes are available for lichen-forming fungi, few surveys of *Starships* in this group have been completed. Yet, given that Pezizomycotina contains several, independently-evolved lichen lineages, it is reasonable to predict the presence of *Starships* in lichen-forming fungi. Indeed, a large-scale screen of *Starships* across ascomycete fungi yielded several predicted *Starships* in the genomes of Lecanoromycete fungi (Supplementary Table 8 in (15)). Here, we analyze four genomes of lichen-forming fungi from the genus *Xanthoria* (Teloschistales, Lecanoromycetes) derived from long-read sequencing data, two of which are newly generated. We describe *Tangerine*, a putative *Starship* element present in *X. parietina*, and report signatures of its cargo horizontal gene transfer (HGT) with a lineage enriched in non-mycobiont fungi that are known to inhabit lichen thalli (Chaetothryiales, Eurotiomycetes). Following this, we report that the captain of *Tangerine* is nested within a new phylogenetic family (*sensu* Gluck-Thaler and Vogan (15)) of *Starship* captains that are endemic to Lecanoromycetes. Additionally, we present a wider phylogenetic analysis of captains within lichen-forming and non-mycobiont lichen-associated fungi and report signatures of putative horizontal *Starship* transfers based on incongruencies in captain phylogenies amongst distantly related lichen fungal symbionts.

## Results

### Comparative analysis of long-read *Xanthoria* genomes

We compared long-read genomes of three isolates of *X. parietina* and its close relative, *X. calcicola* (Fig. 1A-B, Fig. S1). The four genomes showed a high level of synteny, with a similar size and number of predicted genes, but differed in repeat content and GC content (Fig. 1C-E, Table S1). For example, in *X. calcicola* repeats accounted for 23% of the nuclear genome, compared to 13% in *X. parietina*. In all examined genomes, long terminal repeat (LTRs) retrotransposons accounted for a large portion of repeat content (Fig. S2, Table S2). By contrast, long interspersed nuclear elements (LINEs) are absent in the *X. calcicola* genome, yet account for nearly a half of the repeat content in *X. parietina* (Fig. 1D). In all the *Xanthoria* genomes analyzed, repeat elements were not randomly dispersed but instead organized into specific regions of the genome (Fig. S3, Fig. S4). In addition to high repeat density, these regions also had low GC content and showed signs of repeat-induced point mutation (RIP), and were therefore classified as large RIP-Affected Regions (LRARs, defined as genomic regions >4 kb affected by RIP) (20). RIP is a fungal-specific mutation-based mechanism of genome defense against TEs which acts during the sexual cycle (21). RIP appears to have had a more prominent effect on the frequently-asexual *X. calcicola* compared to the obligately-sexual *X. parietina* (Fig.1D). The LRARs of the former were both larger and more numerous and had a lower GC content compared to the latter. Indeed, LRARs appear responsible for the overall lower GC content of *X. calcicola* genome, as the non-LRAR portion of the genomes showed the same median GC content in all four *Xanthoria* isolates (Fig. 1E).

**Fig. 1.**
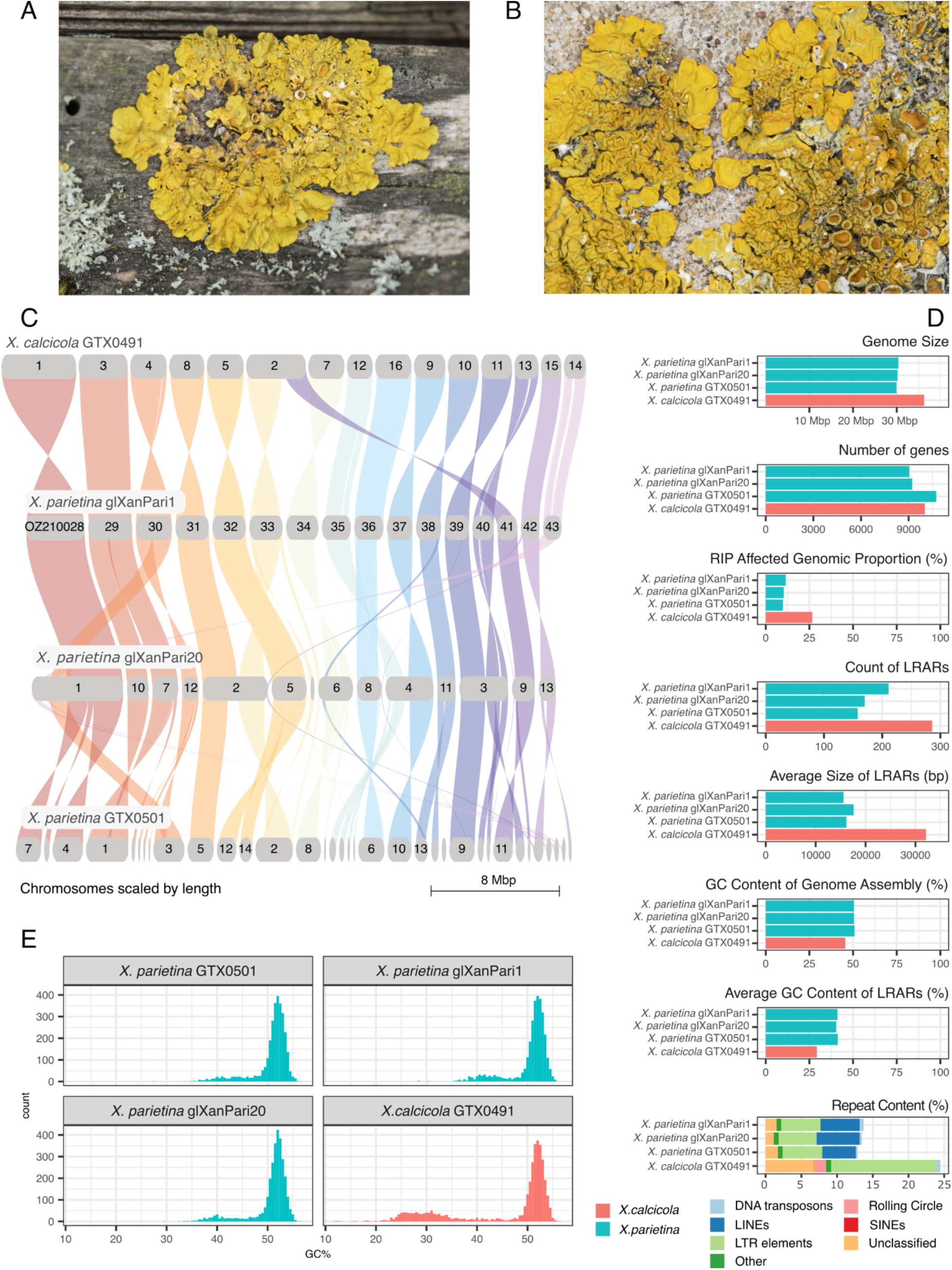
The long-read genomes of *Xanthoria parietina* and *X. calcicola*. **A.** Lichen thallus of *X. parietina* on wood. **B.** Thallus of *X. calcicola* on concrete. **C.** Syntenic map of orthologous regions among three nuclear genomes of *X. parietina* and a genome of *X. calcicola*. Synteny was calculated using GENESPACE based on orthogroup analysis. **D.** Genome statistics for the four *Xanthoria* genomes. Statistics for *X. parietina* genomes are shown as blue bars; for *X. calcicola* as red bars. RIP stands for repeat-induced point mutation; LRAR stands for large RIP-affected region; LINE stands for long interspersed nuclear element; LTR stands for long terminal repeat; SINE stands for short interspersed nuclear element. **E.** Distribution of GC content values across the four *Xanthoria* genomes using non-overlapping windows of 10,000 base pairs.

### Tangerine, a putative Starship element in X. parietina

Using the Starfish pipeline (15), we screened the four *Xanthoria* genomes for the presence of putative *Starship* elements. We identified three elements, each of which contained a tyrR gene and could be aligned to a corresponding empty site in one of the other genomes (Table S3). One of the identified elements was likely a false positive, because it lacked many key characteristics: it was only 17.9 kb long, mapped to an empty site of nearly 2 kb in length, and lacked any genes except for the tyrR. The two remaining putative *Starships*, which we termed *Tangerine,* were found in two *X. parietina* genomes (glXanPari1 and glXanPari20). They belong to the same navis (based on the amino acid similarity of the tyrR captain) and haplotype (based on k-mer based similarity across the entire element sequence).

*Tangerine* architecture is consistent with that of a *Starship* transposon (Fig. 2). Its size (102 kb in *X. parietina* glXanPari1 and 85 kb in *X. parietina* glXanPari20) is close to the median *Starship* length (15). The captain is located 1,390 bp downstream of the 5’ end of the element; it is the first gene in the 5’ direction (Figure 2A). In addition to the tyrR-encoding captain, *Tangerine* contains typical auxiliary genes encoding a protein with a DUF3723 domain, a patatin-like phosphatase (PLP), and a NOD-like receptor (NLR) (15). The boundaries of *Tangerine* were cleanly defined by alignment to the empty site in the genome of *X. calcicola* (Fig. 2A-B). The ends of *Tangerine* contain both 4-5 bp direct repeats and 10-12 bp terminal inverted repeats (Fig. 2C), which notably were separated by 14 bp in the 5’ end of the element and 28 bp in the 3’ end of the element, more than in other reported *Starships* (13).

**Fig. 2.**
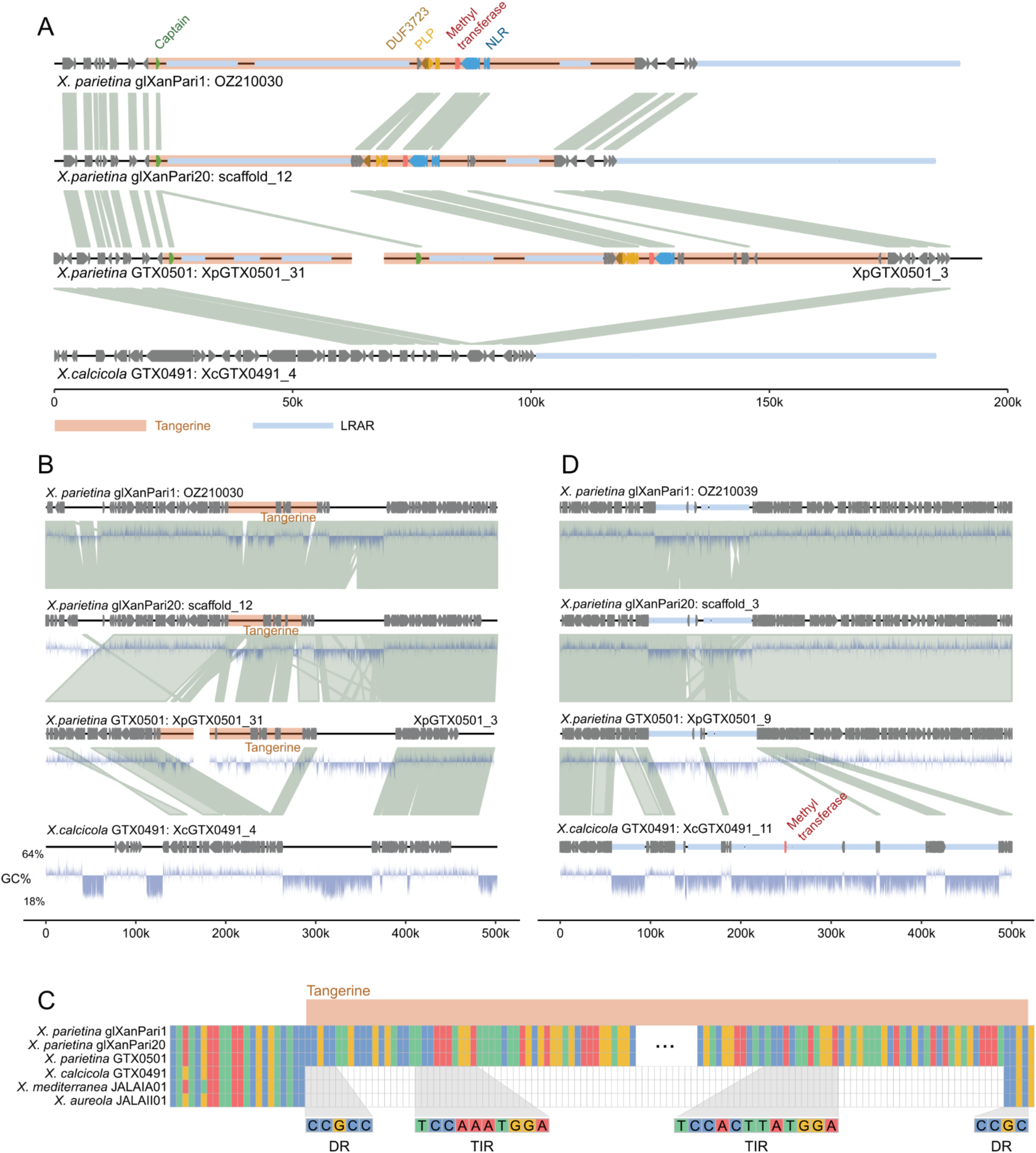
*Tangerine*, a *Starship* element in the genomes of *Xanthoria* fungi. **A.** *Tangerine* (highlighted in orange) in three *X. parietina* genomes aligned to the empty site in *X. calcicola*. In the genome assembly of *X. parietina* GTX0501, *Tangerine* is split between two contigs. Genes are shown as arrows; the captain, accessory and cargo genes are highlighted in different colors. The links between genomic fragments show genes belonging to the same orthogroup. Large Repeat induced point mutation Affected Regions (LRARs) are shown as greyblue bars. PLP stands for Patatin-like phospholipase, NLR stands for NOD-like receptor. **B.** Alignment of *Tangerine* and 200 kb flanking regions. Links between chromosome fragments indicate nucmer nucleotide alignments with >90% sequence identity and >3000 bp in length. The lower track shows the GC-content. **C.** Alignment of *Tangerine* and its flanking regions to the empty site in *X. calcicola* and the short-read genomes of *X. aureola* and *X. mediterranea*. Putative direct repeats (DR) and tandem inverted repeats (TIR) are shown as predicted by the automated Starfish annotation. **D.** Putative degraded copy of *Tangerine* in the genome of *X. calcicola* aligned to the corresponding regions of the three *X. parietina* genomes. Methyltransferase gene is highlighted in red. Blue highlights correspond to LRARs.

*Tangerine* was present in all examined *X. parietina* genomes at the same position. At first, *Tangerine* appeared to be missing in *X. parietina* GTX0501, however upon closer examination, we found the element present in the same region but split between two contigs (Fig. 2A-B), consistent with this genome being more fragmented than others (Fig. 1C). The splitting resulted in two copies of the captain being identified in this assembly, both of them were assigned to the same orthologous group as the captains from *X. parietina* glXanPari1 and glXanPari20 (Table S4). While short-read genomic assemblies are generally too fragmented to allow for *Starship* profiling (15), we confirmed that the 5’ end of *Tangerine* is located in the same region in all eight short-read genomes of *X. parietina* from our earlier metagenomics study (2) (Fig. S5). Since we only detected *Tangerine* at a single genomic locus, we do not have an indication of whether it is still transpositionally active, and hence, consider it a putative *Starship*.

*Tangerine* encodes additional cargo genes beyond the captain and auxiliary genes (Table S5). These include seven predicted genes that lack functional annotation, a gene model encoding a protein with an Ankyrin domain, and, notably, a putative methyltransferase annotated as IPR013216 (Methyltransferase type 11) from the S-adenosyl-L-methionine-dependent methyltransferase superfamily. Across the three *Tangerine* copies in *X. parietina,* predicted proteins from each orthogroup were >95% identical. This was the case for both cargo and auxiliary genes (Fig. S6). The captain sequences were identical across all three genomes.

We used previously produced transcriptomic data from the mycobiont of *X. parietina* and intact *X. parietina* thalli (2) to investigate whether *Tangerine*-associated genes are expressed. Most of the genes, including the captain, were expressed in at least some of the samples, with the NLR and methyltransferase showing highest levels of expression in both lichen thalli and the fungus cultured separately in the lab (Fig. S7, Fig. S8). Out of seven individual lichens from which metatranscriptomic data was produced, two had near zero-levels of expression for all *Tangerine*-associated genes, which might indicate that *Tangerine* has been lost in some *X. parietina* individuals.

More than a half of *Tangerine* (63% in *X. parietina* glXanPari1 and 54% in *X. parietina* glXanPari20) consists of smaller TEs, which is consistent with previous reports of TEs nesting within *Starships* (14, 22) *Tangerine*-associated TEs are affected by RIP and are mostly assigned to LTR/Ty3 and LINE/Tad1 from the same families that dominate repeat content in the rest of the genome (Fig. 2A, Table S6). In addition, a 47-61 kb LRAR is located 13 kb downstream of the 3’ end of *Tangerine* in the *X. parietina* genomes and can be also detected in *X. calcicola*, where it is present 13 kb downstream of the empty site. This LRAR corresponds to the 100 kb largely unaligned region downstream of *Tangerine* (Fig. 2B), in which only small fragments, mostly corresponding to LTR/Ty3 TEs, could be aligned between *X. parietina* and *X. calcicola* (Fig. S9). Compared to *X. parietina*, the LRAR in *X. calcicola* has a lower GC content and a larger portion of repeats that could not be assigned to known superfamilies (32% compared to 0.4% in *X. parietina*; Fig. 2B, Fig. S9), perhaps owing to sequence modification caused by intensive RIP.

To test for the possibility of misassembly, we examined read alignments to the boundaries of *Tangerine* in *X. parietina* glXanPari1 and to the empty site in *X. calcicola* (Fig. S10). No likely misassembly was detected. At the same time, we detected regions with an uneven depth of coverage within *Tangerine* (Fig. S10). Using the read alignments, we annotated structural variants (SVs) within the element and in 200 kb flanking regions. This analysis yielded 16 SVs, of which 10 were present within *Tangerine* (Table S7). The majority of the SV were indels present in GC-poor regions lacking genes, downstream of the captain or upstream of the 3’ end of *Tangerine*. Two notable exceptions are a break point 907 bp downstream of the captain and an insertion within the gene model XANPAOZ2100_002164-T1 (DUF3723), which appears to represent an intron. An identical intron is present in the corresponding gene in *X. parietina* glXanPari20, but is absent from *X. parietina* GTX0501.

### *Tangerine* may be present in other *Xanthoria* species at different locations

To test whether the *Tangerine* element can be located anywhere else in the genome of *X. calcicola,* we searched all *Tangerine*-associated genes in the predicted proteome, genome assembly, and raw reads generated from the *X. calcicola* GTX0491 sample from this study. This search yielded one hit: the amino-acid sequence of predicted protein XANCAGTX0491_008091-T1 was 95% identical to the methyltransferase from *Tangerine* cargo (Fig. S6). This similarity was in line with the similarity of most ortholog pairs in *X. calcicola* and *X. parietina* (Fig. S11). The gene was located in a different contig compared to the empty site described above and was flanked by two LRARs, 61.5 and 45 kb in length (Fig. 2D). Compared with the corresponding region in the three *X. parietina* genomes, *X. calcicola* sequence is inflated by numerous repeat-rich LRARs, which resulted in patchy alignment between the genomes of the two species. It is possible that a copy of *Tangerine* was present in *X. calcicola* at this site, and that it has been degraded by RIP (as has been shown for other *Starships* (13)) until the methyltransferase gene was left as the only remaining “fossil”. Alternatively, the methyltransferase may have been captured and transposed by a different mobile element.

In short-read genome assemblies of *X. mediterranea* and *X. aureola* (23), we found the same empty site as in *X. calcicola* (Fig. 2C, Fig. S12A). Similarly to *X. calcicola*, in the assembly of *X. aureola* we identified only the *Tangerine*-associated methyltransferase, perhaps reflecting a similar lost element, or possibly the original fungal locus picked up as cargo in the *Tangerine*. In *X. mediterranea* we identified the captain, all the auxiliary genes, and the methyltransferase, but these were split between three contigs (Fig. S12B) suggesting that a more intact copy of *Tangerine* may be present in this species at a different genomic locus than that of *X. parietina*.

### The *Tangerine* captain defines a Lecanoromycetes-specific family of *Starships*

Initial HMM-based profiling by Starfish assigned the *Tangerine* captain to clade 1 of the *Starship* tyrR tree, but did not confidently assign it to any of the previously described families. This contrasted with other putative tyrRs detected in *Xanthoria* genomes, many of which were assigned to families such as Phoenix and Galactica (Table S4). To investigate the *Tangerine* captains further, we aligned their sequences to the sequences of 1,222 representative *Starship* tyrRs from Gluck-Thaler & Vogan (15), and reconstructed a phylogeny. In the resulting tree, the *Tangerine* captain was recovered together with three tyrRs from fungi in the same class Lecanoromycetes (Fig. S13).

To determine whether the *Tangerine* clade met the standards of being considered a new *Starship* family, we supplemented the dataset with 1,508 tyrRs (dereplicated into 494 representative sequences using amino acid based sequence clustering) that we identified in a larger set of 554 genomes from lichen-forming and lichen-associated fungi (Table S8). In the reconstructed phylogeny, the *Tangerine* clade sat at the basal part of clade 1 and contained exclusively tyrRs from Lecanoromycetes, most of them lichen-forming (Fig. 3A-B, Fig. S14, Table S9). The clade met the threshold to be recognized as a new *Starship* family (Tangerine-family), as described by Gluck-Thaler & Vogan (15) (>20 representative captains forming a monophyletic clade with >80% SH-ALRT and >95% ultrafast bootstrap support). We performed an additional search against an unfiltered set of captains from 1,649 diverse fungal genomes from Gluck-Thaler & Vogan (15), but failed to recover any Tangerine-family sequences from non-Lecanoromycete fungi.

**Fig. 3.**
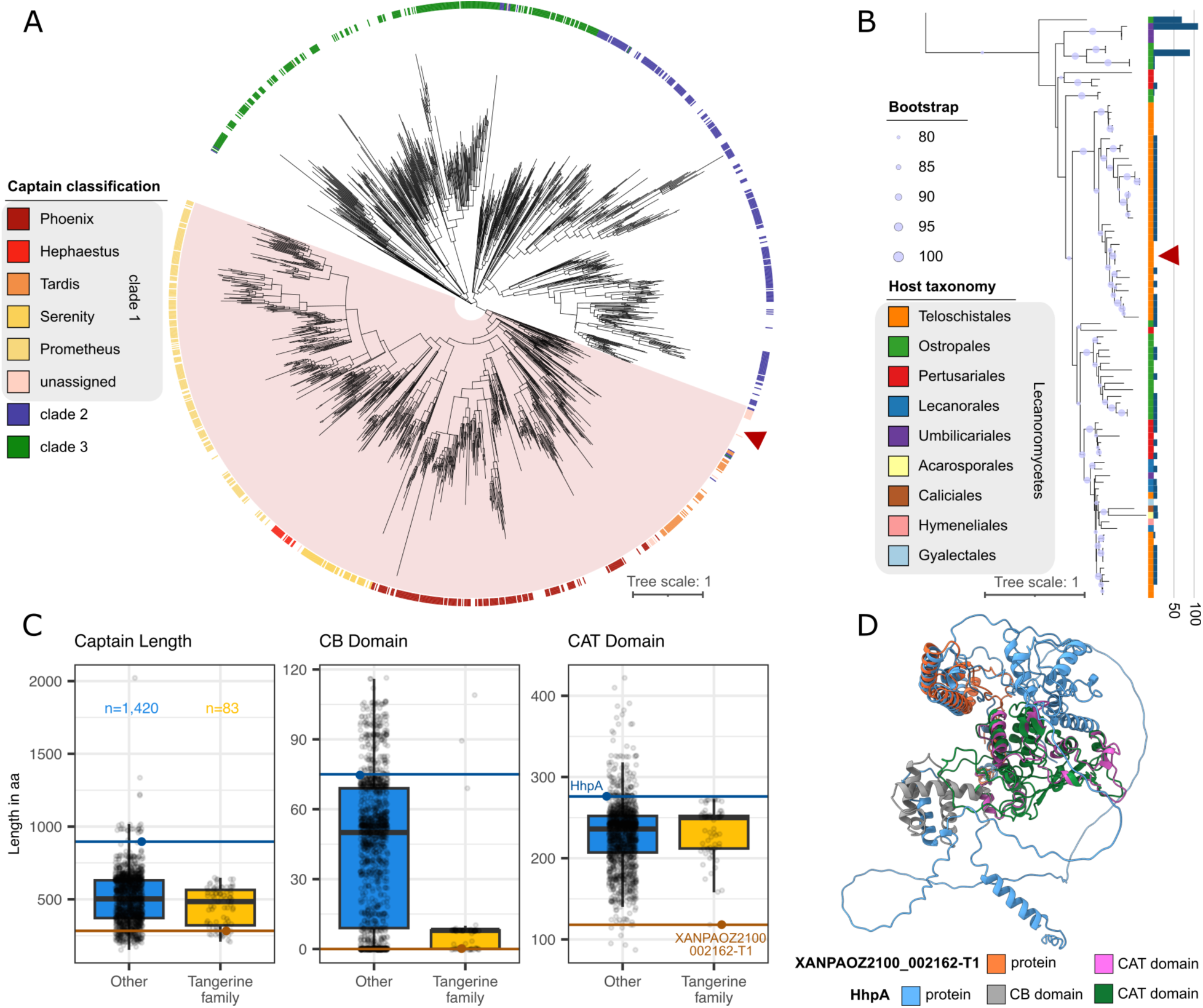
*Tangerine* in the context of the *Starship* captain phylogeny. **A.** Sequence-based *Starship* captain tree with the Tangerine-family indicated by the red arrow. Predicted captains from lichen-associated fungi were clustered into representative sequences and aligned with 1,222 representative *Starship* captains; the alignment was used to generate a maximum-likelihood phylogeny with 100 bootstrap replicates. Clades with <70 bootstrap support are collapsed. The full tree in Newick format with bootstrap values, together with the sequence alignment can be found in the supplemental data files (FigShare). The outer color track shows captain family assignments; clade 1 is highlighted as a pink block. **B.** Magnified clade of the Tangerine-family showing the full set of captains (both representative and not), annotated based on the taxonomy of their host. The *Tangerine* captain from *X. parietina* is indicated with the arrow. Bootstrap support shown next to the nodes. The bars to the right of the tree show the length of the CB (core-binding) domains for each captain. The full tree with taxa labeled is in Fig. S14. **C.** Distribution of full captain, CB domain, and catalytic (CAT) domain lengths (aa) between the Tangerine-family (n=83) and other *Starship* captains (n=1,420). The Tangerine-family set includes the complete set of unclustered captains from the family. The other families are represented by captains annotated *de novo* in genomes of lichen-associated fungi and a subset of captain representatives from Gluck-Thaler and Vogan (15). The *Tangerine* captain from *X. parietina* (XANPAOZ2100_002162-T1) is highlighted in orange, and HhpA captain from *Hephaestus* in blue. **D.** Structurally aligned, AlphaFold3-generated structural predictions of the *Tangerine* captain from *X. parietina* (XANPAOZ2100_002162-T1) and HhpA. Different colors indicate conserved domains: CB and CAT in HhpA and CAT in XANPAOZ2100_002162-T1.

While the median length of captains from the Tangerine-family did not differ from the other *Starship* families, they had a markedly shorter core-binding (CB) domain, one of the characteristic DNA binding domains of tyrosine recombinases (Fig. 3C). Over a third (34.9%, n=29) lacked a CB domain entirely, and an additional 53% (n=44) had a putative CB domain only eight amino acids long and with no similarity to the CB domain of the well-characterized *Hephaestus* captain, HhpA (Fig. S15), suggesting that the CB domain was lost in these captains as well. To confirm that the absence of a CB domain was not an artifact of annotation, we searched HhpA against the nucleotide sequence of the *Tangerine* element, which yielded no hit for the CB domain. In contrast, the larger Tangerine*-*family captains tended to retain the catalytic (CAT) domain (Fig. 3C), and all conserved active sites within it (Table S10). Notably, however, the captain of the *Tangerine* element from *X. parietina* had a short sequence of 283 amino acids and lacked two of the conserved canonical active sites out of six (sites 1 and 2).

To validate the results of the sequence-based analyses, we predicted and analyzed the *Tangerine* captain structure. A predicted structure of the captain monomer was generated via AlphaFold 3 (Fig. S16) and searched against the Protein Data Bank (PDB), which yielded no significant (*E*-value <0.001) results (Table S11). However, when this *Tangerine* captain was structurally aligned to and compared against the predicted structure of the HhpA captain from the well studied *Hephaestus Starship* (Fig. 3D, Fig. S17A), strong structural similarity was observed between the CAT domains with a root mean square deviation (RMSD) value of 1.89 Å across the 134 atomic pairs. Three of the six characterised active sites within HhpA had corresponding residues within the *Tangerine* captain (Fig. S17B). No region of the *Tangerine* captain could be identified to be corresponding to the CB domain found in the *Hephaestus* captain (Fig. 3D, Fig. S17A). To further explore the lack of a CB domain within the Tangerine-family captains, HhpA was compared to an additional Tangerine-family captain from *Caloplaca sp.* (Fig. S17C), which contained eight residues of the CB domain. However, the truncated putative CB domain from the *Caloplaca sp.* captain corresponds structurally to part of the region in between the CB and CAT domains on HhpA (Fig. S17D), providing another line of evidence suggesting the CB domain may be lost in most of the Tangerine-family captains.

To investigate the structural relationships between the Tangerine-family and other *Starship* captains, we generated structural phylogenies using Foldtree based on various distance matrices derived from protein-protein alignments (24). The phylogeny based on the Foldtree metric, which combines both local structure and sequence information, reveals the Tangerine-family captains form a distinct clade (Fig. S18). However, both structure-only phylogenies, based either on local distance difference tests (LDDT) or template modelling score (TM), had large inconsistencies with the sequence-based phylogeny. Only the LDDT-based phylogeny formed a Tangerine-family clade, which did not include some members of the Tangerine-family, including the captain from the *Tangerine* element in *X. parietina* (Fig. S18).

### *Tangerine*-associated genes show signatures of horizontal gene transfer

*Starships* have been implicated as vectors of HGT in fungi (16, 25–27). Within the *Tangerine* element in *X. parietina*, one auxiliary gene, the NLR, and one cargo gene, the methyltransferase, appear to have been horizontally transferred. The two genes neighbour each other and are separated by only 350 bp. When searched against the NCBI nr database, these genes returned numerous high-quality hits (>95% identity over >95% sequence) to immediate relatives of *Xanthoria* from the order Teloschistales and to distantly-related Chaetothyriales fungi (Eurotiomycetes). Crucially, no hits to Lecanoromycete fungi other than from Teloschistales were obtained (Fig. S19). To rule out the possibility that these genes were missing from the NCBI nr database due to variable genome annotation, we searched the NCBI nt database but failed to obtain a non-Teloschistales Lecanoromycete hit of a better quality than hits to Chaetothyriales (Table S12). The Alien Index (AI) (28) for these two genes was 461 and 284 respectively, which is well above 45, the threshold for likely HGT (Fig. 4A, Table S13). This pattern was unique to the NLR and methyltransferase and was not detected in other *Tangerine*-associated genes, flanking genes in the vicinity of *Tangerine*, or 10 randomly selected genes from the genomic background as controls (Fig. S20, Fig. S21).

**Fig. 4.**
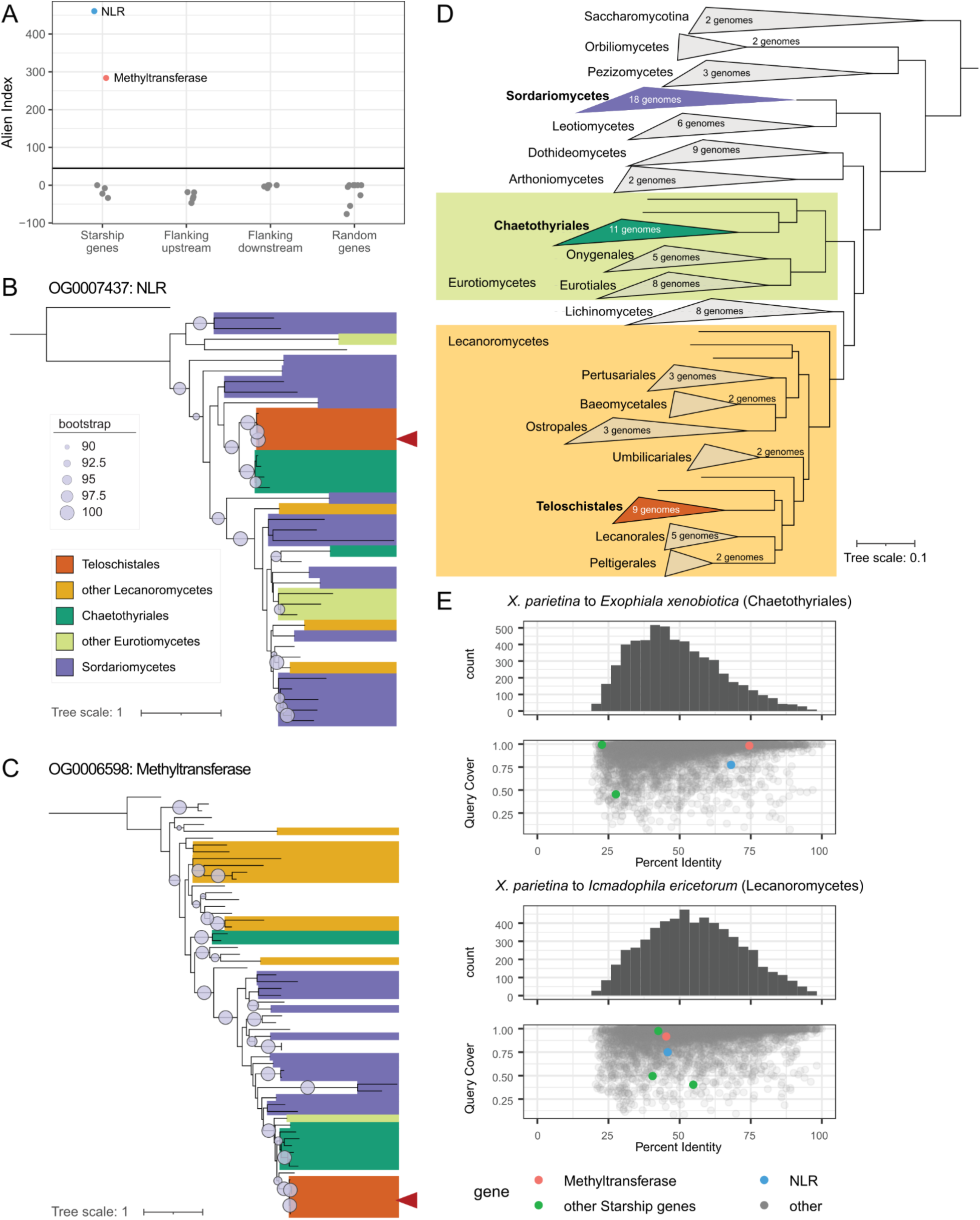
Putative horizontal gene transfer (HGT) of *Tangerine*-associated genes. **A.** Alien Index (AI) of *Tangerine*-associated genes, gene models in the immediate vicinity of *Tangerine*, and 10 randomly selected gene models from *X. parietina* glXanPari1. AI is defined as ln(best E-value of non-lecanoromycete hit + 1e^-200^) - ln(best E-value of non-Teloschistales lecanoromycete hit + 1e^-200^). The horizontal line corresponds to the AI of 45, which is considered the threshold for likely HGT (28). **B-C.** Gene tree for the orthogroup containing *Tangerine*-associated NOD-like receptor (NLR, B) and methyltransferase (C). To obtain the trees, we performed orthogroup analysis on 107 ascomycete genomes (Table S14). For the selected orthogroups, we computed maximum-likelihood phylogeny. The tree is colored based on the taxonomic origin of sequences; bootstrap values are shown for the clades with >90 support. The sequences from *X. parietina* glXanPari1 are indicated with arrows. Full trees with taxa labeled are in Fig. S25, Fig. S26. **D.** Species tree generated by Orthofinder for the 107 ascomycete genomes. The tree is rooted using two Saccharomycotina genomes. **E.** Cross-mapping of the predicted proteome of *X. parietina* glXanPari1 to *Exophiala xenobiotica* and *Icmadophila ericetorum*. For each *X. parietina* glXanPari1 predicted protein, we identified the best match in the other predicted proteome. The top panel shows the distribution of percentage identity values across all matches; the bottom panel shows how the matches were positioned in relation to percentage identity and query cover. NLR, methyltransferase, and other *Tangerine*-associated genes are highlighted in color.

To further examine the evolutionary histories of the putative HGT candidates, we ran an orthogroup analysis on 107 Ascomycete genomes (Table S14). Gene trees for the NLR and methyltransferase-containing orthogroups were discordant compared to the species tree (Fig. 4B-D, Fig. S22, Fig. S23). For both genes, a well-supported clade of genes from Teloschistales is closely related to Chaetothyriales genes. In the case of the NLR, the Teloschistales clade is sister to the Chaetothyriales clade. In the case of the methyltransferase, Teloschistales clade is nested within Chaetothyriales. Notably, the same methyltransferase clade contains genes from two additional distantly related lichen-forming fungi: *Chaenotheca gracillima* (Coniocybales, Lichinomycetes; protein similarity 76% over 99% coverage) and *Endocarpon pussilum* (Verrucariales, Eurotiomycetes; protein similarity 73% over 98% coverage). In both gene trees, the clade containing Teloschistales and Chaetothyriales genes is nested within a larger clade consisting primarily of genes from Sordariomycete fungi. Again, the discordance between gene trees and the species tree was observed only for the NLR and methyltransferase, and not for other *Tangerine-*associated genes or genes in its vicinity (Fig. S24, Fig. S25). The two orthogroups including the NLR and methyltransferase were present in 21 genomes included in the analysis. Excluding *Xanthoria*, in eight genomes the two genes are located in the vicinity of each other, and only in three genomes – two from Chaetothyriales and one from Sordariales – did the ‘tail-to-tail’ gene configuration match that observed in *Tangerine* (Fig. S26).

Finally, we cross-mapped the proteomes of *X. parietina* with *Icmadophila ericetorum*, a fungus from the same class of Lecanoromycetes, and *Exophiala xenobiotica*, a Chaetothyriales fungus whose genome encoded homologs of the key *Tangerine*-associated genes (Fig. 4E). Both the NLR and methyltransferase had higher percentage identity between *X. parietina* and *E. xenobiotica* compared to the more closely-related pairing of *X. parietina* and *I. ericetorum*.

### Other *Starships* may be present in *Xanthoria* genomes

In addition to the captain of the *Tangerine* elements, the four *Xanthoria* genomes encoded 16 other putative tyrR genes which grouped into nine orthologous gene groups i.e. naves (plural, from singular ‘navis’, latin for ‘ship’) (15) (Table S4). While none of them was associated with a predicted element with clearly defined boundaries, several were positioned within 100 kb of genes encoding PLP or NLR proteins, or other *Starship* auxiliary genes. As more high quality *Xanthoria* become genomes available, we may be able to validate whether any of these non-*Tangerine* tyrRs are associated with bonafide *Starship* elements.

### Three captain tyrR clades show signatures of horizontal *Starship* transfer amongst distantly-related lichen fungal symbionts

We used the newly constructed phylogenetic tree of tyrR captains to identify other instances of possible HGT between distantly-related lichen-forming and non-mycobiont lichen-associated fungi, this time based on incongruencies between the captain tree and the species tree. In the captain tree, we located three clades of interest (Clades A, B, and C), in which captains from different classes of lichen-associated fungi clustered together in a way that is discordant with the species tree (Fig. 5A-C, Fig. S27, Fig. S28, Fig. S29, Table S15). Clade A is nested within the Galactica-family and consists primarily of captains from lichen-forming Lecanoromycetes and the Eurotiomycete lichen *Endocarpon pusillum*, as well as non-mycobiont lichen-associated Chaetothryiales (Fig. 5A,D). This clade specifically shows a high degree of discordance with the species tree suggesting multiple potential HGT events. Clade B is positioned within the Tardis-family and suggests HGT events from Eurotiomycetes to Lecanoromycetes lichen-forming fungi (Fig. 5B,D). Since Eurotiomycetes and Lecanoromycetes classes are quite closely related (Fig. 4D), this result warrants more caution. At the same time, since the clade overwhelmingly consists of sequences from lichen-associated fungi with only two sequences from non-lichen Eurotiomycetes present, HGT within the context of a lichen symbiosis remains a plausible explanation. Clade C was recovered within the Prometheus-family and presents evidence for possible HGT between distantly related lichen-forming fungi from the classes Lecanoromycetes and Arthoniomycetes (Fig. 5C-D). Given that Arthoniomycetes is a sister clade to Dothideomycetes and is even more distantly related to Lecanoromycetes and Eurotiomycetes (Fig. 4D), HGT appears as the most likely explanation. While this additional captain-based evidence further indicates that *Starship*-mediated HGT may occur between fungi united by their lichen-associated lifestyles, further studies based on more contiguous genomes are necessary to validate the described instances of HGT.

**Fig. 5.**
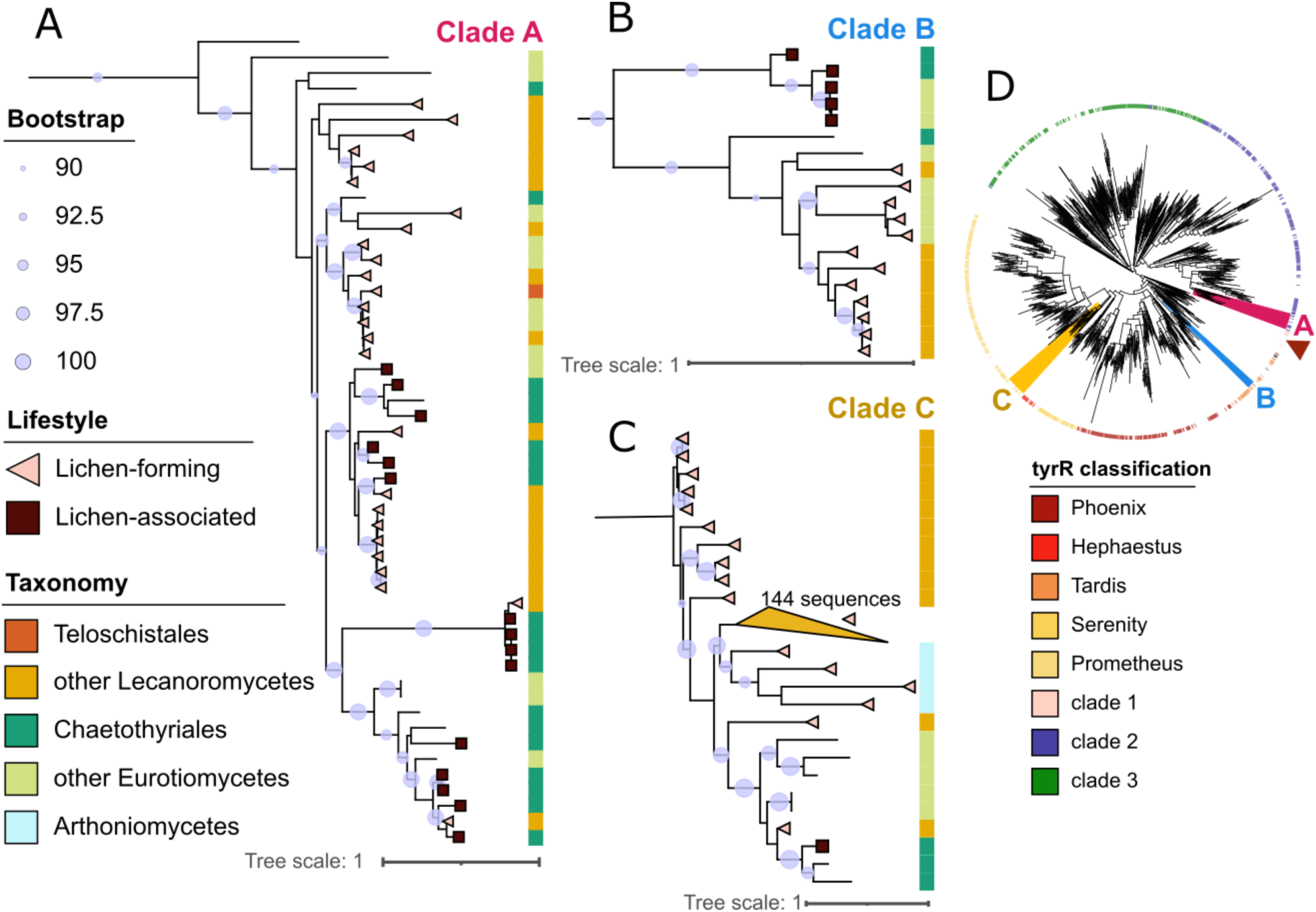
Clades of *Starship* tyrR captains exhibiting signatures of horizontal gene transfer (HGT) between distantly-related lichen symbionts. A-C. Phylogenetic clades of interest. Sequences from all tyrR captains (both representative and not) within clades from the larger captain phylogeny that showed signatures of HGT in lichen fungi were re-aligned, and the alignment was used to recreate more refined clade-specific maximum-likelihood phylogenies. Bootstrap values are shown for branches with >90 support. The symbols at the ends of branches indicate lifestyle: tyrRs from lichen-forming fungi (also known as mycobionts) are shown in pink triangles, tyrRs from non mycobiont lichen-associated (also known as lichenicolous or endolichenic) fungi are shown as brown squares. Sequences from non-lichen fungi lack annotation. The color strip is colored according to the taxonomic group. Full annotated trees are shown in Fig. S27, Fig. S28, Fig. S29. Information on the sequences from Clades A-C is shown in Table S15. **D.** Clades A-C in the context of the larger tyrR tree. The color strip shows the family and clade assignments. The Tangerine-clade is indicated by the red arrow.

## Discussion

Since their recent discovery, *Starships* have been implicated in the evolution of multiple features crucial to the lifestyle of their fungal hosts, including specialized metabolism, virulence, and abiotic stress relief (29–33). We therefore set out to investigate whether the lichen lifestyle may also be shaped by *Starship* activity. Here, we provide the first detailed report of a putative *Starship* element in the genome of a lichen-forming fungus. We named the element *Tangerine* in reference to the yellow-orange color of *Xanthoria* lichens and after a spacecraft briefly featured in one of Douglas Adams’s novels (34). *Tangerine* possesses all the hallmark characteristics of a *Starship*, including its large size, boundaries defined by direct repeats four base pairs long, and characteristic gene content, including a tyrosine recombinase captain gene, auxiliary genes, and cargo.

We found the *Tangerine* element in the genomes of *Xanthoria parietina*, commonly known as the sunburst lichen. Across the genus *Xanthoria*, we had access to several genomes of sufficient quality to allow *Starship* detection, which is rarely the case for lichen-forming fungi, enabling this analysis to be carried out. Whether the *Tangerine* element in *Xanthoria* is still mobile or whether it has lost the ability to transpose is not yet clear. In all of the *X. parietina* genomes analyzed, including short-read genomes used for additional screening, *Tangerine* was always found in the same genomic region, suggesting it no longer is mobile. In the other three surveyed *Xanthoria* species, however, this region contained an empty site. *Tangerine* may be present elsewhere in the genome of *X. mediterranea*, but its exact location cannot be determined due to a lack of long contiguous regions in the genome assembly. In the long-read genome of *X. calcicola*, only a potential degraded copy of *Tangerine* could be detected, again positioned at a distinct site in the genome. When considered together, this evidence suggests that if *Tangerine* is not currently active, it may have been transposing as recently as the time of *Xanthoria* diversification. Regardless of whether *Tangerine* is still mobile or not, the *Tangerine* captain, auxiliary genes, and cargo are expressed in lichen thalli, which may indicate their encoded products have continued to play some biological role in the fungus.

By annotating captains in a larger set of lichen-associated fungi we revealed *Tangerine* to be the first representative of a new family of *Starships* endemic to the Lecanoromycetes. The CB domain is truncated in this family, which may indicate altered or loss of DNA binding activity. Yet, loss of the CB domain does not necessarily imply the loss of recombination activity, as several bacterial tyrosine recombinases with no CB domain can nonetheless recognize and bind DNA and carry out recombination (35). Notably, the *Tangerine* captain from *X. parietina—*the only member of the Tangerine-family whose exon boundaries could be validated using RNA data —has a truncated CAT domain in addition to the loss of CB domain, raising further questions about its functionality. However, the overall conservation of the *Tangerine* captain’s predicted structure with the *Hephaestus* HhpA captain, coupled with the fact that it has maintained its position at the 5’ terminus of a putative *Starship*, suggests it still plays some role in the biology of *Starship* transposons. Future research will clarify whether Tangerine-family captains are capable of transposition and/or some other function.

*Starship* cargo genes often perform functions beneficial to their hosts and directly involved in their lifestyle, such as effector genes in plant pathogenic fungi. Since little is known about the machinery of the lichen symbiosis, our ability to make such inferences regarding the cargo of the *Tangerine* element is limited. However, one gene, the methyltransferase (XANPAOZ2100_002166-T1 in *X. parietina* isolate glXanPari1) does stand out. This gene was the only *Tangerine-*associated gene also detected in the genomes of *X. calcicola* and *X. aureola*. In *X. calcicola*, it appears within a potential degraded copy of *Tangerine* as the only gene surrounded by large RIP-affected regions. Such conservation amidst extensive RIP is unusual and might suggest that the gene is being maintained by selection because it may be beneficial to these lichen-forming fungal species. This might be analogous to the plant pathogen *Parastagonospora nodorum*, which carries an intact and active virulence factor *ToxA* in a degraded copy of the *Horizon Starship* (*32*). The gene was assigned to the S-adenosyl-L-methionine (SAM)-dependent methyltransferase superfamily. SAM-dependent methyltransferases are ubiquitous and act on a wide range of substrates (36). Among other functions, SAM-dependent methyltransferases are involved in secondary metabolism, including synthesis of polyketides, a class of metabolites prevalent in lichens (37, 38). It is plausible that this gene could be involved in the biosynthesis of natural products. The role of this gene and its potential *Starship*-mediated transfer therefore warrants further investigation.

Two *Tangerine-*associated genes, including the SAM-dependent methyltransferase, appear to have been horizontally transferred from other fungal taxa. The exact origin of the genes is difficult to pinpoint, but our phylogenetic analysis suggests origins either in the Chaetothyriales (Eurotiomycetes) or in the Sordariomycetes. Both of these groups harbour *Starships* (39), which suggests the possibility of *Starship*-mediated HGT; however, we cannot rule out that *Starships* acquired these genes after they had been horizontally transferred through some unknown mechanism. Chaetothyriales fungi consist primarily of black yeasts with diverse ecologies (40), including notably non-mycobiont lichen-associated fungi, which are present in lichen thalli in small quantities (41, 42). Indeed, one of the Chaetothyriales fungi harbouring a highly similar methyltransferase gene, *Knufia peltigerae* (Fig. S23), has been described from a lichen thallus (43). Moreover, in a recent analysis of *X. parietina* metagenomes, we reported several Chaetothyriales lineages present sporadically in *X. parietina* thalli (2). Since lichen mycobionts can co-exist with Chaetothyriales in the context of the symbiotic lichen thallus, HGT between the two groups is therefore plausible. Notably, one of the two genes, the methyltransferase also appears to have been transferred not only to the order Teloschitales, including *Xanthoria*, but also to the genomes of additional two lichen-forming fungi from different classes of Ascomycota, which might suggest a link to the lichen lifestyle. This transfer is not likely to be recent, as the methyltransferases from these fungi share only about 70% sequence similarity.

*Tangerine-*mediated HGT is not the only instance of possible HGT between distantly related lichen-associated fungi that we found. By examining the phylogeny of *Starship* captains, we identified three clades within the broader tyrR captain phylogeny with signatures of HGT amongst lichen-forming Lecanoromycetes, lichen-forming Eurotiomycetes, lichen-forming Arthoniomycetes, and lichen-associated Chaetothryiales (Eurotiomycetes). Host relatedness has been hypothesized to drive horizontal *Starship* transfer (39), but additional ecological factors might be at play. For example, these HGT events may have been driven by spatial proximity: either between two lichens living in the same microhabitat, or between a lichen-forming fungus and a lichen-associated (‘endolichenic’) fungus within the same lichen thallus. Our ability to explore these possibilities was limited by the fact that the majority of the fungal genomes available for this analysis were assembled from short-read data—the assembly quality permitted annotation of captains but not the associated *Starship* elements. Future analyses using long-read genomes will stress-test the validity of the potential HGT events and allow us to explore potential links between host ecology and *Starships*. The presence or absence of ecologically-relevant cargo genes within these *Starships* may shed light onto the drivers of lichen-mediated HGT.

Our dataset also allowed us to compare genomes of two closely related lichen-forming fungi that differ in the frequency of sexual reproduction. While largely similar, the genomes showed stark differences in repeat content, with the often-asexual *X. calcicola* having both higher repeat content and more pronounced RIP. This difference might indeed be linked to the difference in frequency of sexual cycle in *X. calcicola*. TEs might accumulate in the asexual propagation phase and then be efficiently RIPed during more rare crossing events, whereas frequent meiosis in *X. parietina* may prevent accumulation of repeats and therefore lead to fewer RIPed regions. In lichen-forming fungi, closely related species often differ in presence or absence of sexual reproduction (44). For these fungi, the evolutionary consequences of sex are larger than for non-symbiotic fungi. In addition to effects on genome evolution and recombination, for lichen fungi the presence or absence of sex also affects their transmission strategy. Sexually reproducing species have to undergo ‘resynthesis’ and re-establish the lichen symbiosis following ascospore germination, while asexual lichen-forming fungi typically co-transmit with their symbionts in larger vegetative propagules containing both fungus and alga/cyanobacterium (1). Future research will clarify the trade-offs associated with different lichen reproductive strategies, both in connection to genome evolution and the evolution of symbiosis.

Lichen biology remains largely unknown and mysterious, in large part due to the numerous methodological challenges lichens pose as study organisms. Our discovery of the *Tangerine* element in *Xanthoria*, the *Tangerine-*family of *Starships* in Lecanoromycetes, and signatures of *Starship* HGT amongst distantly-related lichen symbions, however, raises the possibility that these unusual transposons may play a role in lichen biology. As long-read genome sequencing becomes more accessible, more high-quality genomes of lichen fungi will undoubtedly be generated from lichen metagenome studies allowing further evaluation of our hypothesis that *Starships* contribute to the evolution of the lichen lifestyle. The presence of *Tangerine* in *Xanthoria* genomes and signatures of *Starship*-mediated HGT amongst lichen symbionts provides the basis and rationale for such analyses.

## Methods

### Genome assembly and annotation

To obtain a genome of *Xanthoria calcicola*, we collected a lichen sample from concrete at the Norwich Research Park (Norwich, UK; 52.623133°N, 1.221621°E). The voucher for this specimen is deposited in UC Berkeley University and Jepson Herbaria under ID UC2111092. We extracted metagenomic DNA using a NucleoBond High Molecular Weight DNA Kit. Short fragments were removed with a Circulomics Short Read Eliminator Kit with 25 kb cut-off.

Remaining DNA was sequenced using PromethION Flow Cell FLO-PRO114M to 20.8 Gbp of raw data. We used Dorado v0.2 for basecalling and Flye v2.9-b1780 with flags “overlap 10K, error rate 0.005, no-alt-contigs’’ for assembly. In addition, we produced short-read data from the same DNA extraction using Illumina NovaSeq 6000 platform to produce 2 Gbp of PE150 data. We used short-read data to polish the assembly with Pilon v1.23 (45). To produce the purified genomic assembly of the *X. calcicola* mycobiont, we binned the metagenomic assembly. For that, we first aligned the short reads onto the assembly with Bowtie v2.4.1 (46) and binned the assembly with MetaBAT v2.15 (47). To assess the quality of the resulting genome assembly, we used BUSCO v4.0.6 (48) with the ascomycota_odb10 database. To confirm the taxonomic identity of the fungus, we located the ITS sequence, combined it with published sequences of *Xanthoria* (Table S16), and aligned the set of sequences using MAFFT v7.271 (49). We trimmed the alignment to remove positions absent in over 90% of sequences using trimAl v1.2 (50) and reconstructed the phylogeny using IQ-TREE v2.2.2.2 (51) using 10,000 rapid bootstraps (Figure S1).

To obtain the *X. parietina* genome glXanPari20 (isolate SAMEA115359166), we collected a lichen sample from a tree branch at the Wellcome Genome Campus (Hinxton UK; 52.07891°N, 0.184294°E). Samples were prepared for DNA extraction and Hi-C sequencing using a CryoPrep for flash frozen tissue pulverization. DNA extraction was performed using a potassium chloride extraction buffer with chloroform as highlighted in Park et al. (52). DNA extracts were sheared with a MegaRuptor at a speed of 130. Sheared DNA was SPRI cleaned up (0.88v/v concentration) with RNase to degrade any remaining RNA material. Final DNA extracts were submitted for PacBio HiFi Sequencing on the Revio platform through Scientific Operations at the Wellcome Sanger Institute. Pulverized lichen tissue was submitted for Hi-C sequencing using the Arima V2 kit on the Illumina NovaSeq X through Scientific Operations at the Wellcome Sanger Institute. PacBio HiFi reads were assembled with metaMDBG v0.3 (53). To determine assembly coverage, reads were mapped back to the assembly using minimap2 v2.23 (54). Contigs were binned with MetaBAT2 v2.12.1 (47) and bins were assessed for quality using EukCC v2.1.2 (55) and BUSCO v5.7.0 (48). Final scaffolds were generated using the Arima Hi-C mapping pipeline (https://github.com/ArimaGenomics/mapping_pipeline/blob/master/Arima_Mapping_UserGuide_A160156_v03.pdf) and yahs v1.2 (56).

Four genomes were analyzed: *X. parietina* isolate GTX0501 from Tagirdzhanova et al. (2) (PRJEB78723; genome accession GCA_964263705.1), *X. parietina* glXanPari1 from the isolate SAMEA111342678 (PRJEB83421; genome accession GCA_964656405.1), and the two genomes described above. The genomes were annotated using Funannotate v1.8.15 (57). For *X. parietina* GTX0501 we re-used the annotations from Tagirdzhanova et al. (2) the remaining genomes were annotated in the same way as follows. The genomes were repeat-masked using stand-alone RepeatMasker v4.1.2 (https://www.repeatmasker.org/). For *X. calcicola* and *X. parietina* GTX0501 we used repeat libraries created with RepeatModeler v2.0.3 (58) with the - LTRStruct flag; for the other *X. parietina* isolates, we re-used the library produced for GTX0501. Next, we ran the ‘funannotate predict’ module, which used Genemark-ES v4.62 (59), Augustus v3.3.2 (60), CodingQuarry v2.0 (61), GlimmerHMM v3.0.4 (62) for *de novo* gene prediction, created consensus models with EVidence Modeler v1.1.1 (63), and annotated tRNA with tRNAscan-SE v2.0.9 (64). To produce functional annotations, we used the ‘funannotate annotate’ module and following databases: PFAM v35.0 (65), UniProtDB v2023_01 (66), MEROPS v12.0 (67), dbCAN v11.0 (68), and BUSCO ascomycota_odb10 (48). Separately, we ran InterProScan v5.42-78.0(69) and the EMapper webserver (http://eggnog-mapper.embl.de/) (70). For family-level annotation of repeat elements, we used EarlGrey v5.1.1 (71) with the Dfam v3.8 (72). To annotate genome regions affected by repeat-induced point mutations we used RIPper (20). To visualize GC-content of the genomes, we used seqkit v0.12.0 (73). To visualize global synteny, we used GENESPACE v1.3.1 (74).

### *Starship* detection

To detect potential *Starships*, we used Starfish v1.0.0 (15) following the tutorial (https://github.com/egluckthaler/starfish/wiki/Step-by-step-tutorial). Briefly, we ran ‘starfish annotate’ and ‘starfish consolidate’ modules to detect tyrosine recombinases. Next, we ran ‘starfish insert’ and ‘starfish summarize’ to identify insertion sites and element boundaries, and ‘starfish dereplicate’ to assign putative elements to genomic regions. To visualize the elements we used the ‘starfish locus-viz’ module and gggenomes v1.0.1 (75). The region alignments were produced with MUMmer v4.0.0 (76). To assess read coverage of the *Starship* element in *X. parietina* glXanPari1, we mapped raw data (SRA: ERR13660094) on the genome assembly using Minimap2 v2.24-41122 (54) and visualized the alignment with IGV (77). To align the element ends, we used MAFFT v7.271 (49). To locate *Tangerine* in *X. calcicola* and *X. parietina* GTX0501, we searched for sequences similar to *Tangerine*-associated genes in the genomic assemblies and raw metagenomic reads using blast+-2.9.0 (78). We also searched for homologues of *Tangerine*-associated genes in the short-read metagenome-assembled genomes of *X. parietina* and visualized the gene expression data from our earlier study (2). To locate *Tangerine* and the flanking genes in the short-read genomes of *Xanthoria* from Llewellyn et al. (23) (JALAII01 and JALAIA01), we used results of the orthogroup analysis (see description of the HGT analysis). To compare proteins encoded within *Tangerine* in different genomes, we used blast+-2.9.0 (78). To confirm the transcript annotation for the *Tangerine*-associated genes, we aligned two metatranscriptomic libraries from *X. parietina* (SRA: ERR4235132 and ERR4235133) to each *X. parietina* genome using STAR v2.5.4b (79) and cross-referenced the exon boundaries with the RNA alignments. This way, we corrected exon boundaries in the captain genes in *X. parietina* glXanPari1 (XANPAOZ2100_002162-T1) and glXanPari20 (XANPARI20_008752-T1) (Fig. S30). Both the initial annotation and the RNA-corrected version differed from the annotation produced for this gene model by Starfish (15), which relied on MetaEuk (80) for de novo gene prediction. Starfish predicted an extra exon at the N-terminus of the protein, which however was not supported by the RNA data and not included in the final gene model (Fig. S30).

To further test that the CB domain and a portion of CAT domain is genuinely truncated in the *Tangerine* captains, we attempted to find additional exons that would cover the ‘missing’ portion of the *Tangerine* captain. We used a tBLASTn search of the *Hephaestus* captain HhpA against the nucleotide sequence of the *Tangerine* element, which yielded no additional hit outside of the *Tangerine* captain gene model.

### Captain characterization

To place the *Tangerine* captain in the families of other *Starships*, we combined its sequence with the 1222 selected representative sequences of tyrRs from Gluck-Thaler and Vogan (15) and Crypton sequences from fungi (AAO92638.2 and KAE8209120.1), aligned them with MAFFT v7.271 (49) using the E-INS-i method. We trimmed the alignment to remove positions absent in over 70% of sequences using trimAl v1.2 (50) and, following Gluck-Thaler and Vogan (15), extracted 512 first positions from the trimmed alignment and reconstructed the phylogeny using IQ-TREE v2.2.2.2 (51) using with automated model selection, 1000 SH-ALRT, iterations and 1000 ultrafast rapid bootstraps.

Following the observation of the *Tangerine-*like captains forming a Lecanoromycetes-specific clade, we tested whether this clade forms a new lichen-associated *Starship* family. We used ‘starfish annotate’ module from Starfish v1.1.0 (15) to annotate captains in genomes of lichen-associated fungi from Lecanoromycetes, Eurotiomycetes, Dothideomycetes, and Arthionomycetes from NCBI (Table S17). A total of 4,128 putative captains were identified. We added the captain sequences from *X. parietina* and *X.calcicola* to this dataset and clustered it to produce putative active representative captain sequences, following Gluck-Thaler and Vogan

(15). First, captain sequences shorter than 203 amino acids (n=1,481) were removed. Next, we removed putative pseudogenes defined as sequences missing >1 of the six conserved active sites in tyrRs (n=1,139), excluding the *Tangerine* element captains from *X. parietina*. To annotate active sites, we aligned the lichen captains to representative sequences of tyrRs from Gluck-Thaler and Vogan (15) using MAFFT v7.271 (49) and cross-referenced the alignment with annotated active sites. A large cluster of sequences, including the *Tangerine* captains, had a different column for active sites 3 and 4 (Table S10); filtering for this cluster was done separately. We clustered the remaining lichen captains (n=1,508) into representative sequences using MMseqs v2-13.0 (81). First, we used ‘easy-clust’ to cluster sequences by 90% sequence identity and 80% overlap with the longest seed sequence of the cluster. The resulting representative sequences were clustered again, by 50% sequence identity and 80% overlap. We then aligned these representative captains (n=494) with the 1,222 representative sequences of tyrRs from Gluck-Thaler and Vogan (15) and 3 outgroup tyrR sequences from fungal Crypton DNA transposons (82) using MAFFT v7.271 (49) with the E-INS-i method. Columns with ≥90% gaps were removed using ClipKIT v2.7.0 (83).

We constructed tyrR phylogeny using the CB and CAT domains plus 50 aa flanking sites, which we annotated by cross-referencing the crypton sequences between our alignments and the trimmed alignment from Gluck-Thaler and Vogan (15). We then trimmed these sites in AliView (84), and constructed phylogeny using IQ-TREE v2.2.2.2 (51) with the automated model selection, 1000 SH-ALRT iterations, and 1000 ultrafast rapid bootstraps. The tree was rooted with the crypton tyrRs using PhyKIT v2.1.2 (85). Finally, we confirmed that the Tangerine-family is specific to Lecanoromycetes, by searching the unfiltered set of 10,771 captains annotated by Gluck-Thaler and Vogan (15) and clustering these sequences with the Tangerine-family captains by 50% sequence identity and 80% overlap using MMseqs v2-13.0 (81), which failed to recover any non-Lecanoromycetes captains.

The structural-based phylogenies were generated from a set of 696 predicted models using Foldtree (24). Initially a set of 745 captains was chosen combining a manually selected representative group of *Starship* captains from clade 1 from Gluck-Thaler and Vogan (15), the *X. parietina Tangerine* captain XANPAGTX0501_001716-T1, and a set of *de-novo* annotated representative captains from lichen-associated fungi. The set of 2,684 captains from lichen-associated fungi was filtered to sequences 203–999 (inclusive) residues in length and then clustered by 30% similarity using MMseq2. This resulted in 453 representatives which were then combined into the final set to make 745 captains total (Fig. S31). Structural models of the final set of captains were generated via AlphaFold2 (86), with four failing predictions. The resulting models were then filtered by pLDDT > 50, leaving the 696 models used as input for Foldtree (Table S18). Models for pairwise superimposition were generated via AlphaFold3 (87). For each protein, five predictions were generated and the most confident model was chosen based on pTM and average pLDDT. Model superimposition and RMSD values were generated via the matchmaker function in ChimeraX (88) with the default settings. The PAE plots were visualised using PAE Viewer (89).

### Structural Variant annotation

To call structural variants present in the *Tangerine* in the *X. parietina* glXanPari1 genome, we used Sniffles v2.5.2 (90). First, we mapped raw data (SRA: ERR13660094) on the genome assembly using Minimap2 v2.24-41122 (54). We sorted the read alignment with samtools v1.18 (91), and isolated the portion that corresponded to the putative *Starship* and a 200 kb flanking region on each side. Resulting alignment was analyzed with Sniffles2 using default settings.

### HGT analysis for *Tangerine*-associated genes

To test *Tangerine*-associated genes for HGT signatures, we first searched them against the NCBI nr database using BLASTp web interface (accessed 2025/03/21). As controls, we included five flanking gene models on each side of the *Starship* element, as well as 10 randomly selected gene models. Using the search results for each gene, we calculated the AI following Gladysev et al. (28). Here, we defined AI as ln(best E-value of non-lecanoromycete hit + 1e^-200^) - ln(best E-value of non-Teloschistales lecanoromycete hit + 1e^-200^). To check whether the absence of certain gene models in the NCBI nr database is an artifact of genome annotation, we searched likely-HGT models using tBLASTn web interface for the protein sequences (accessed 2025/05/24) and BLASTn for the transcript sequences (accessed 2025/10/10) against the NCBI core_nt (with no taxonomic restriction) and full nr/nt (restricted to Lecanoromycetes) databases (Table S12).

To further test HGT candidates, we used orthogroup analysis. We combined the four *Xanthoria* genomes with 104 previously published genomes (Table S14) and analyzed them with OrthoFinder v2.5.4(92). Species and gene trees were visualized using iTOL v7(93). We rooted the trees using *Saccharomyces cerevisiae* and *Lipomyces starkeyi* (Saccharomycotina). Using MMseqs v2-13.0 (81), we crossed-mapped predicted proteome of *X. parietina* glXanPari1 and proteomes of two fungi: *Exophiala xenobiotica* (GCA_035930045) and *Icmadophila ericetorum* (GCA_022814295).

For the two orthogroups containing HGT candidates, we produced Maximum-Likelihood phylogenies. For each orthogroup, we aligned the sequences with MAFFT v7.271 (49), trimmed the alignment to remove positions absent in over 80% of sequences using trimAl v1.2 (50), and reconstructed the phylogeny using IQ-TREE v2.2.2.2 (51) using 10,000 rapid bootstraps.

### HGT analysis for *Starship* tyrRs

To find captain clades with signatures of HGT between distantly-related lichen symbionts, we first identified instances of lichen captains from different classes that clustered together during the step of clustering representative captains. Twelve instances of cross-class captain clustering were identified. Of these four were found in clades enriched in lichen-associated captains, defined as clades >70% of sequences from lichen-associated fungi. To further test the HGT hypothesis for each clade, we re-constructed its phylogeney using the full unclustered set of captains. The trees were constructed using IQ-TREE v2.2.2.2 (51) with the automated model selection, 1000 SH-ALRT iterations, and 1000 ultrafast rapid bootstraps. Three clades showed signatures of HGT due to their discordance with the species tree. While the other clade only contained lichen-forming Lecanoromycetes and Eurotiomycetes, the positioning of the captains showed a very weak signature of HGT. This phylogenetic-approach, rather than a sequence similarity approach like an alien index or blast-all analysis, was used to identify more ancient events of HGT with lower sequence similarity of the horizontally-transferred genes.

## Data Availability

Genomic data including raw reads and genome assembly for *X. calcicola* is deposited in ENA (PRJEB96921; genome accession GCA_976602755.1). Raw reads for *X. parietina* glXanPari20 are available at ENA as ERR14947898. Starfish outputs, AlphaFold models, captain sequence alignments, sequence- and structure-based phylogenies, and genomic annotation data are available in FigShare (doi: 10.6084/m9.figshare.30417202). All scripts associated with the analysis are available at GitHub (https://github.com/metalichen/2025-Tangerine-Lichen-Starship).

## Author Contributions

GT, NJT, and EGT planned and designed the study. GT and ESC produced genomic data. GT and NB performed the bioinformatic analyses. AHB performed protein structural modelling and analysis. GT drafted the manuscript. All authors contributed to writing.

## Acknowledgements

This work was supported by grants from The Halpin Family, The Gatsby Charitable Foundation, and the Biotechnology and Biological Sciences Research Council BBS/E/J/000PR9798 to NJT. MCM is supported by a UKRI Future Leaders Fellowship MR/Y01717X/1. EGT and NB are supported by the Office of the Vice Chancellor for Research and Graduate Education at the University of Wisconsin-Madison with funding from the Wisconsin Alumni Research Foundation and the Department of Plant Pathology at the University of Wisconsin-Madison. Work at the Wellcome Sanger Institute was supported by Wellcome award 220540/Z/20/A and by the Aquatic Symbiosis Genomics Project award from the Gordon and Betty Moore Foundation [GBMF8897] to MB. The authors thank the participants and organizers of the Aquatic Symbiosis Genomics project Lichen Hackathon at the Wellcome Sanger Institute for feedback and suggestions on the early stages of this project. We thank Aaron Vogan for advice on tyrR annotation, Haoyu Niu and other staff of the Aquatic Symbiosis Genomics project (94) for early access to the genome of *X. parietina* isolate glXanPari1, Neha Sahu for advice on the horizontal gene transfer analysis, Phil Robinson for the lichen photographs, and Alison MacFadyen for the help with depositing data.

