## Supplementary Figures for "*Tangerine:* a new family of *Starships* from lichen-forming fungi"

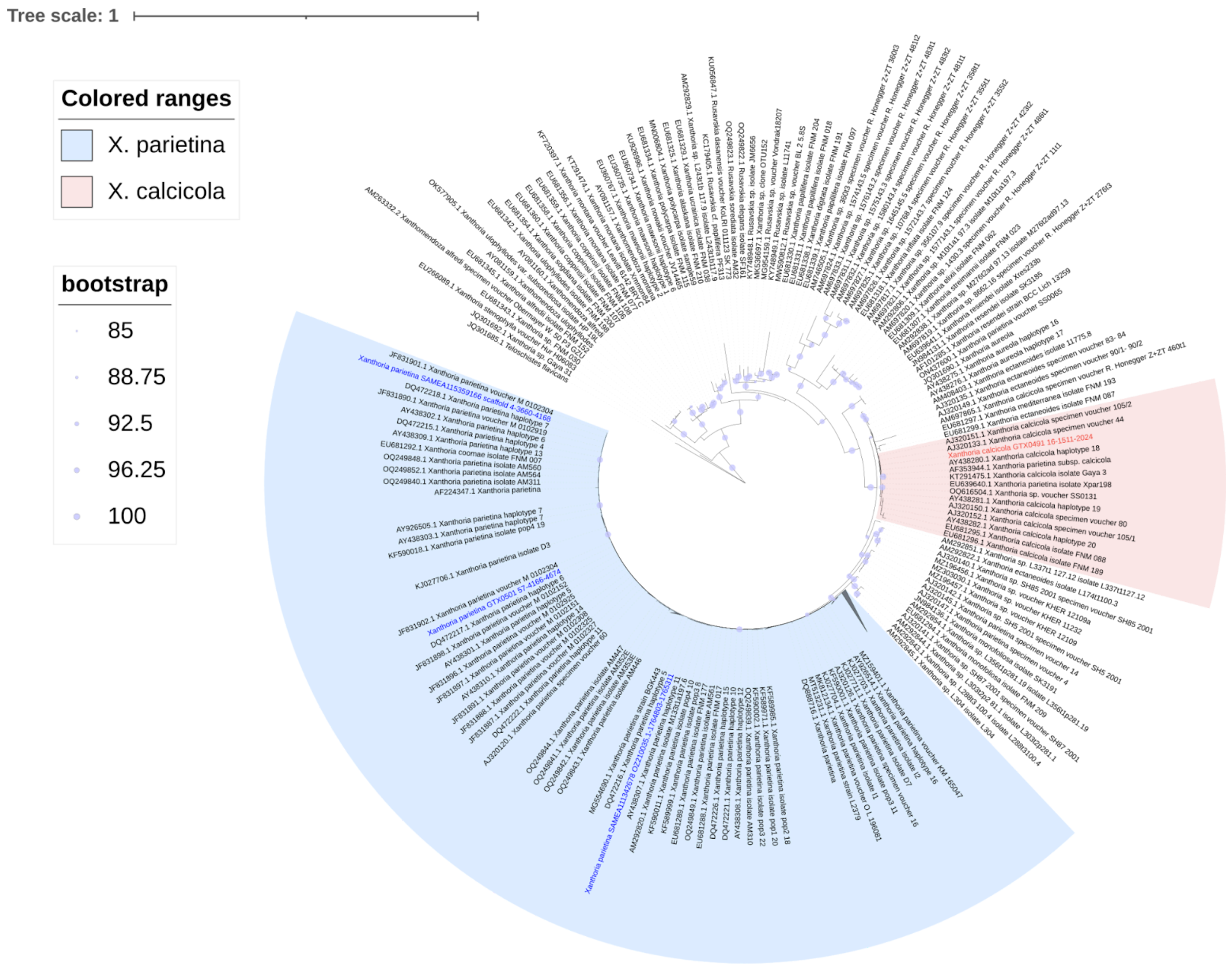


**Fig. S1. Maximum-likelihood phylogeny based on internal transcribed spacer (ITS) sequences.** We combined sequences from literature (Table S16) with ITS sequences extracted from the four *Xanthoria* genomes included in this analysis (highlighted in blue for *X. parietina* and in red for *X. calcicola*). Clades corresponding to *X. parietina* and *X. calcicola* are highlighted.

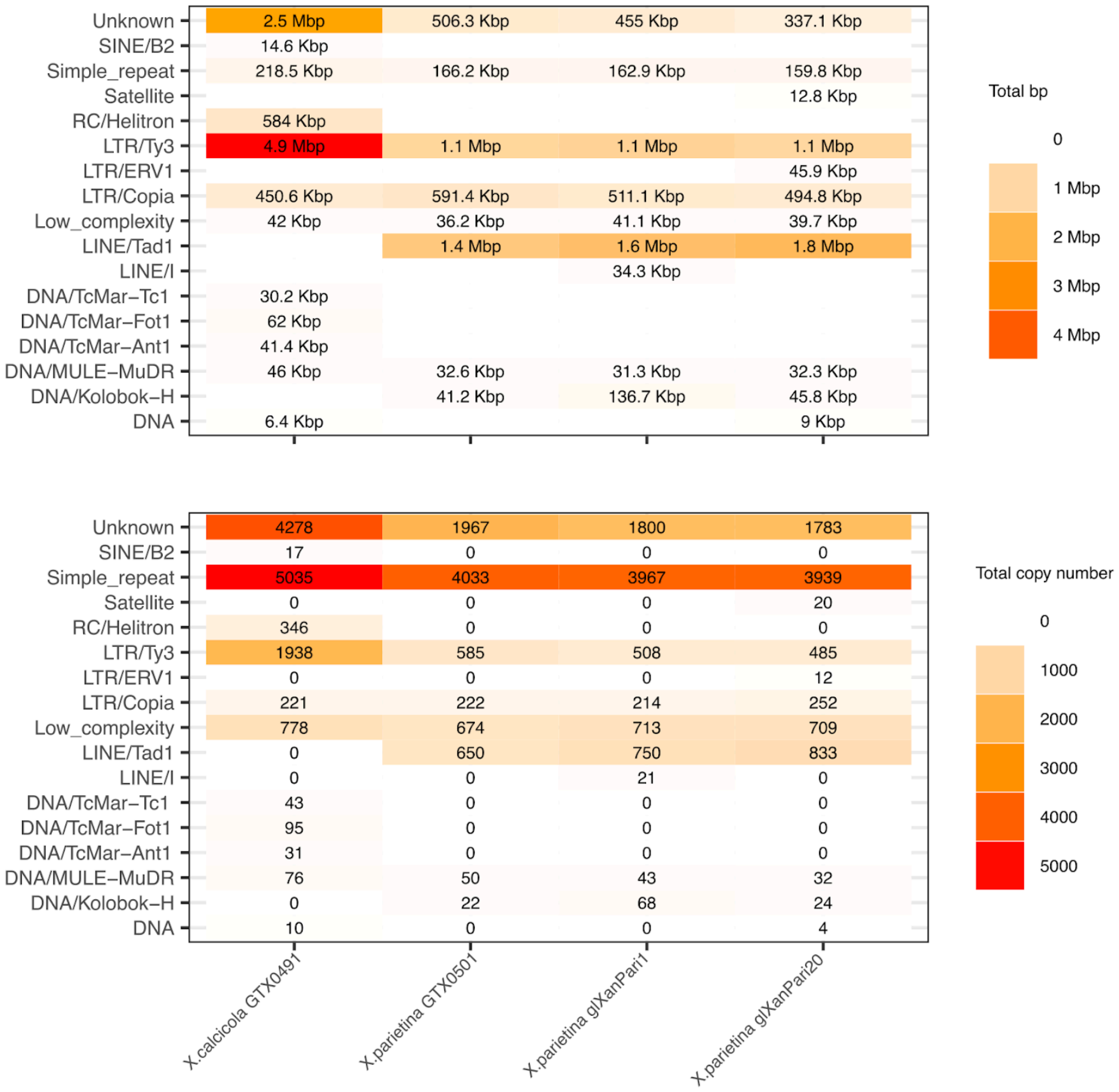


**Fig. S2. Repeat superfamilies present in the four *Xanthoria* genomes.** Repeats were annotated and classified using Earl Grey. The top panel shows total genomic coverage (bp) for each superfamily, summed across all representatives. The bottom panel shows total copy number for each superfamily, summed across all representatives.

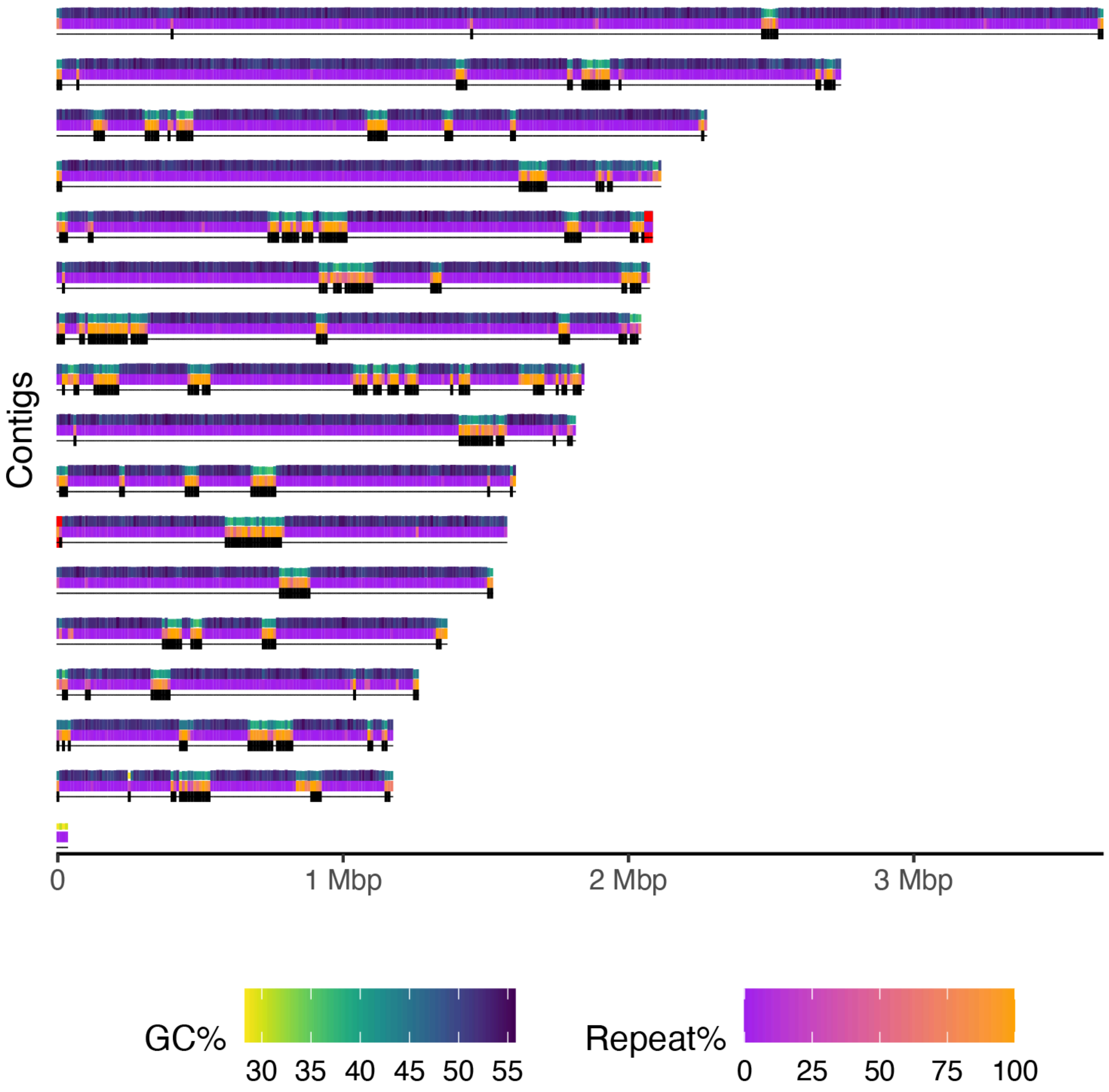


**Fig. S3. The nuclear genome of *X. parietina* isolate glXanPari1.** Each bar represents a contig arranged by the decreasing length, divided into three annotation tracks: GC content, repeat content, and presence of large RIP-affected regions (LRARs). Red bars at the contig ends show telomeric repeats. The x axis shows contig length.

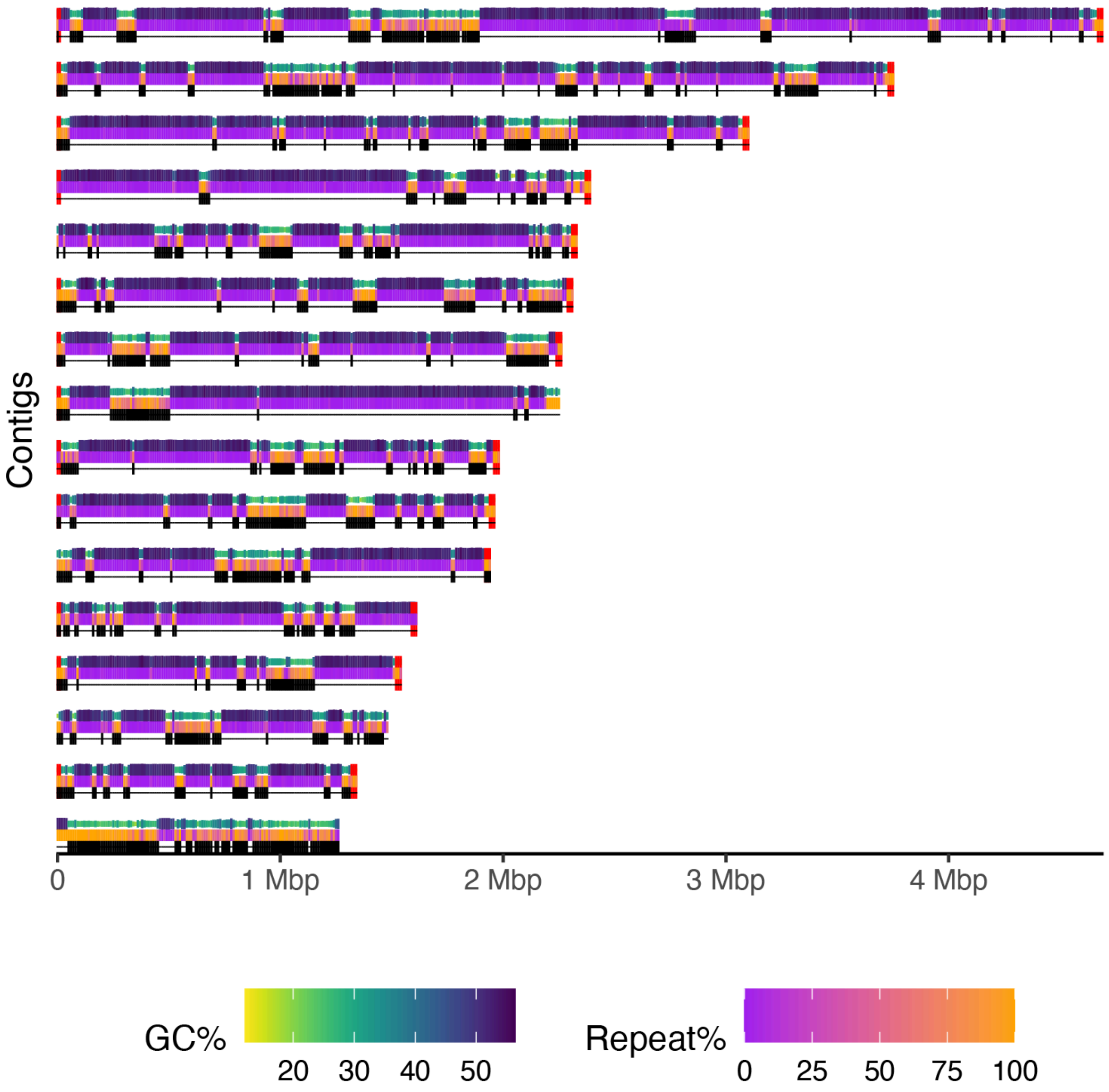


**Fig. S4. The nuclear genome of *X. calcicola* isolate GTX0491.** Each bar represents a contig arranged by the decreasing length, divided into three annotation tracks: GC content, repeat content, and presence of large RIP-affected regions (LRARs). Red bars at the contig ends show telomeric repeats. The x axis shows contig length.

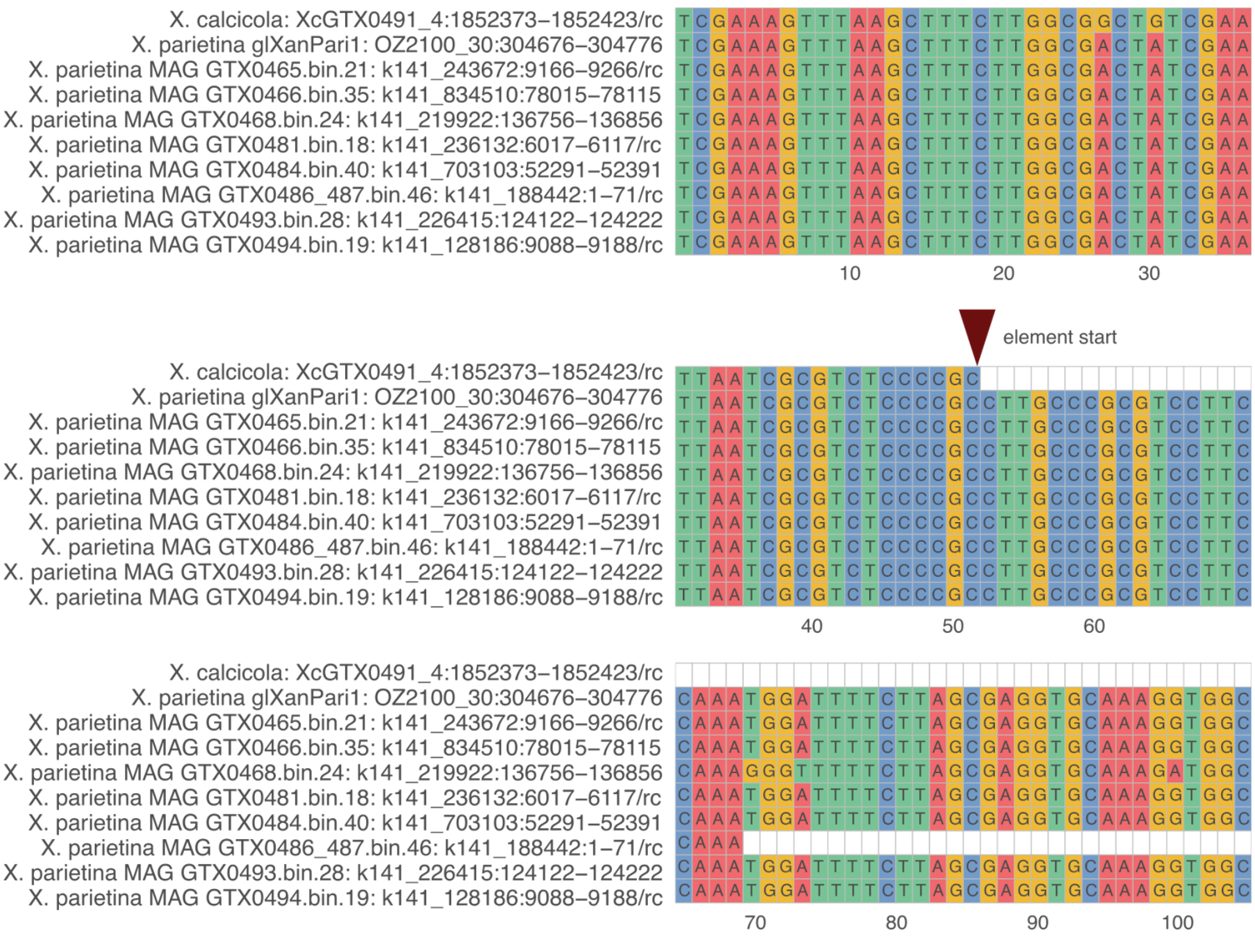


**Fig. S5. Alignment of the first 50 bp of *Tangerine* and 50 bp on the 5' flank in the short-read metagenome-assembled genomes (MAGs) of *X. parietina* from Tagirdzhanova et al. (2025).** The first two sequences represent the empty site in *X. calcicola* and the corresponding region in *X. parietina* glXanPari1 shown for context. The remaining eight sequences show the 5' end of *Tangerine* in the MAGs. In one of the MAGs (GTX0486_487.bin.46), the region of interest was located at the end of the contig, thus explaining the gap in the alignment. In all MAGs except for GTX0486_487.bin.46 and GTX0468.bin.24, the same contig that contained the 5' end of *Tangerine* also encoded a tyrosine recombinase with >99% similarity to the *Tangerine* captain from *X. parietina* glXanPari1 (XANPAOZ2100_002162-T1). In GTX0468.bin.24, the corresponding contig encoded a tyrosine recombinase with 68% similarity to XANPAOZ2100_002162-T1. In GTX0468.bin.24, we failed to get a match to XANPAOZ2100_002162-T1.

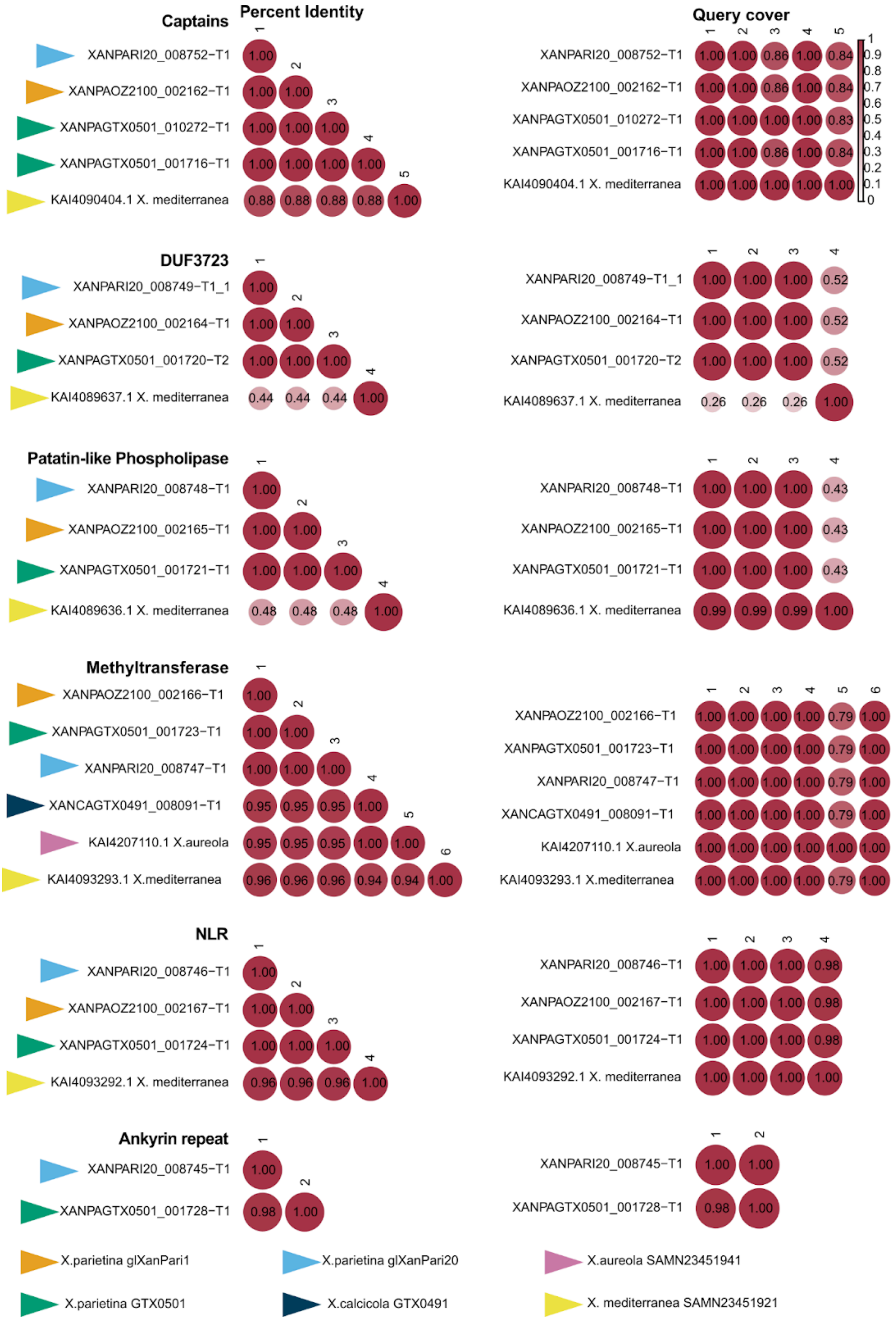
Captains
Percent Identity
Query cover
XANPARI20_008752−T1
XANPAOZ2100_002162−T1
XANPAGTX0501_010272−T1
XANPAGTX0501_001716−T1
KAI4090404.1 X. mediterranea
DUF3723
XANPARI20_008749−T1_1
XANPAOZ2100_002164−T1
XANPAGTX0501_001720−T2
KAI4089637.1 X. mediterranea
Patatin-like Phospholipase
XANPARI20_008748−T1
XANPAOZ2100_002165−T1
XANPAGTX0501_001721−T1
KAI4089636.1 X. mediterranea
Methyltransferase
XANPAOZ2100_002166−T1
XANPAGTX0501_001723−T1
XANPARI20_008747−T1
XANCAGTX0491_008091−T1
KAI4207110.1 X.aureola
KAI4093293.1 X.mediterranea
NLR
XANPARI20_008746−T1
XANPAOZ2100_002167−T1
XANPAGTX0501_001724−T1
KAI4093292.1 X. mediterranea
Ankyrin repeat
XANPARI20_008745−T1
XANPAGTX0501_001728−T1
X.parietina glXanPari1
X.parietina glXanPari20
X.aureola SAMN23451941
X.parietina GTX0501
X.calcicola GTX0491
X. mediterranea SAMN23451921

**Fig. S6. Blast alignments between sequences of proteins encoded within *Tangerine*.** For *X. parietina* glXanPari1 and glXanPari20, only sequences within the *Tangerine* are included; for the other genomes the proteins were selected based on their orthogroup assignments. We aligned sequences from each orthogroup in pair-wise combinations using blast, and visualized percent identity (left column) and query coverage (right column) for each pair. Arrows show which genome the protein originated from. This figure does not show gene models that lacked functional annotation. For the *Tangerine* captain in X. parietina GTX501, we show two gene models, which were present in the genome due to the fragmentation of *Tangerine* onto two contigs.

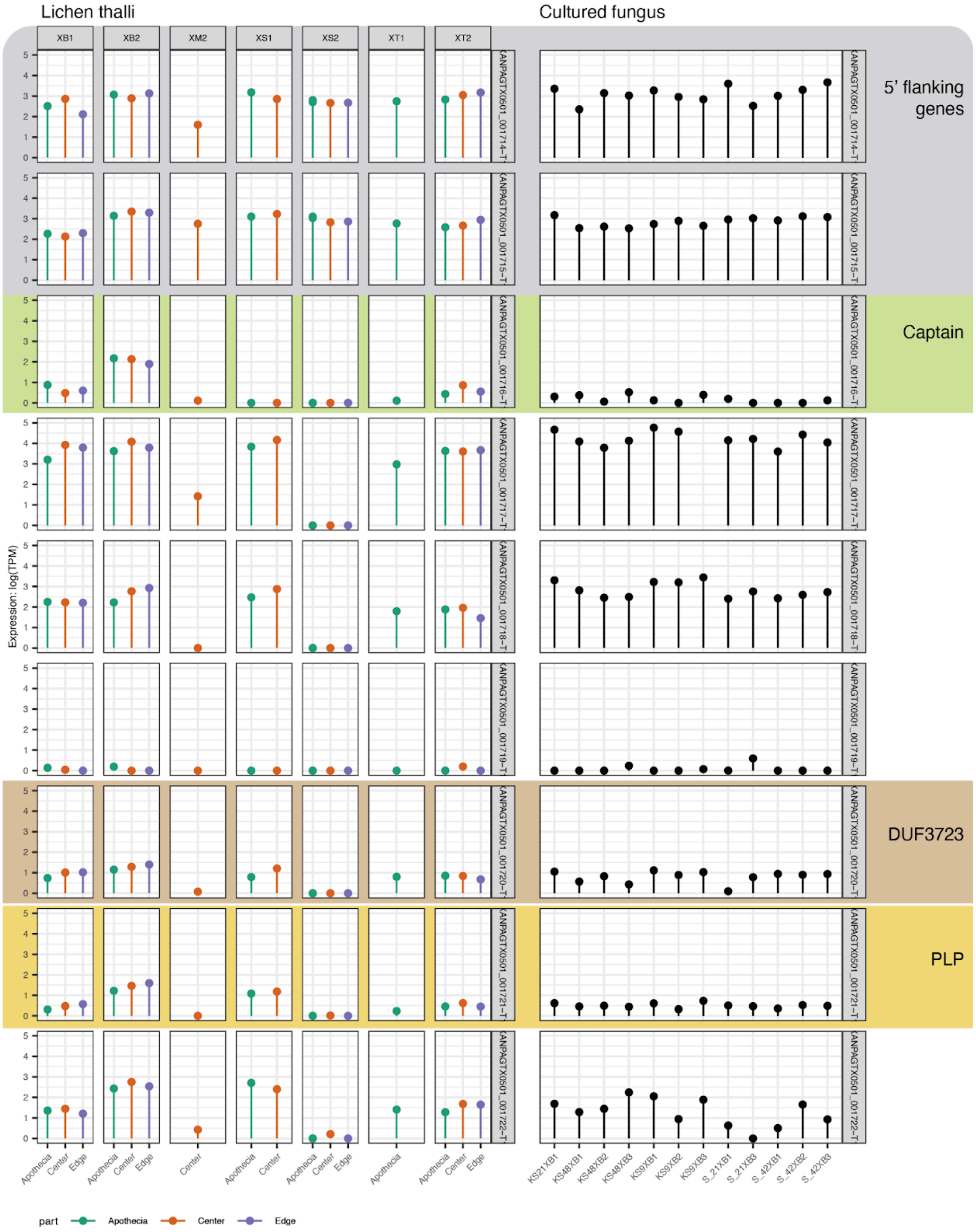


**Fig. S7. Expression of *Tangerine*-associated genes and flanking genes in lichen thalli (left panel) and fungus culture (right panel), part 1.** Lichen thalli samples are grouped based on the thallus and colored based on the developmental stage. Lichen thallus XS2 shows near-zero levels of genes expression for all *Tangerine*-associated genes. The data are taken from Tagirdzhanova et al. (2025; PRJEB38537). TPM stands for transcripts per million, PLP stands for Patatin-like phospholipase, NLR stands for NOD-like receptor.

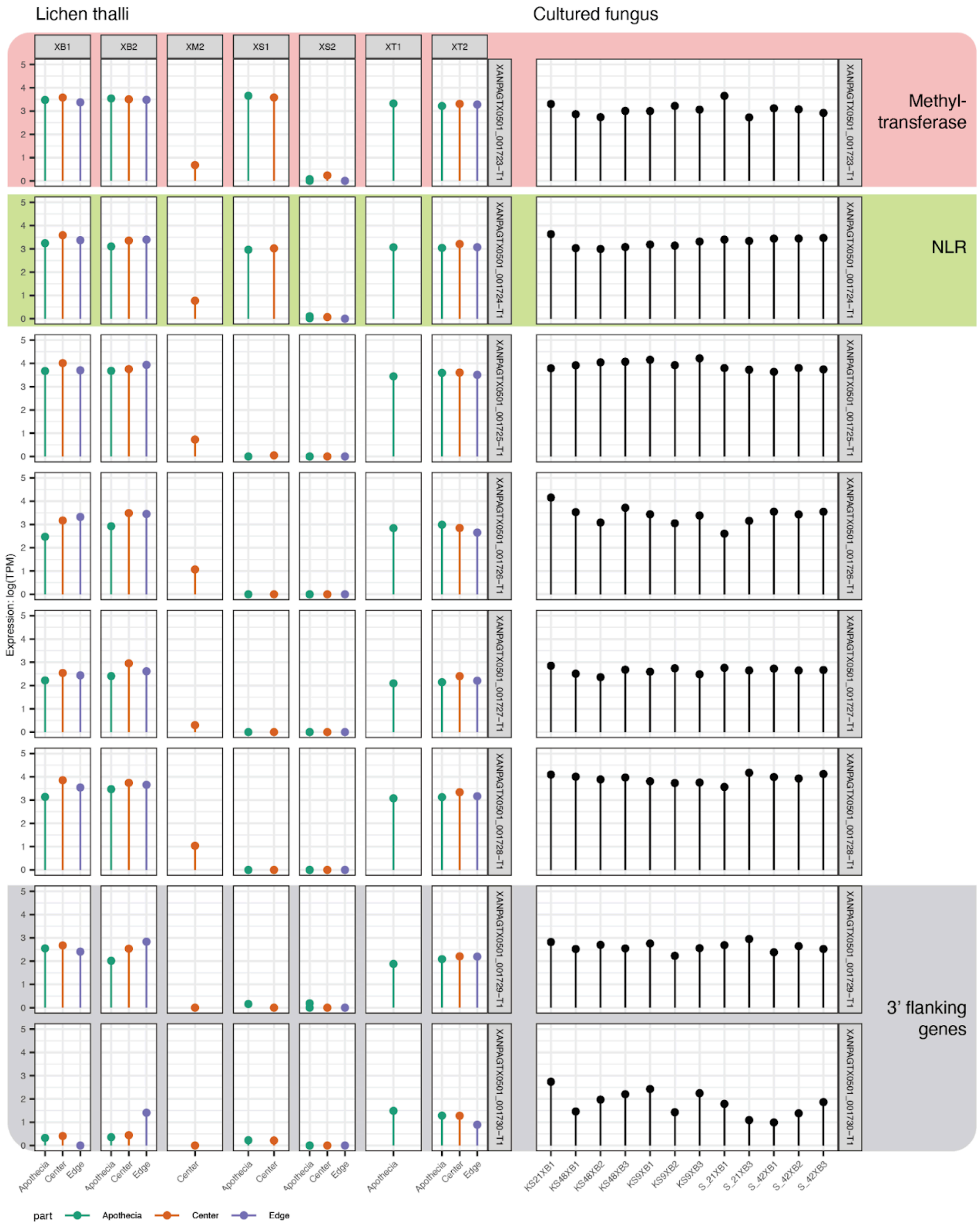


**Fig. S8. Expression of *Tangerine*-associated genes and flanking genes in lichen thalli (left panel) and fungus culture (right panel), part 2.** Lichen thalli samples are grouped based on the thallus and colored based on the developmental stage. Lichen thallus XS2 shows near-zero levels of genes expression for all *Tangerine*-associated genes. The data are taken from Tagirdzhanova et al. (2025; PRJEB38537). TPM stands for transcripts per million, PLP stands for Patatin-like phospholipase, NLR stands for NOD-like receptor.

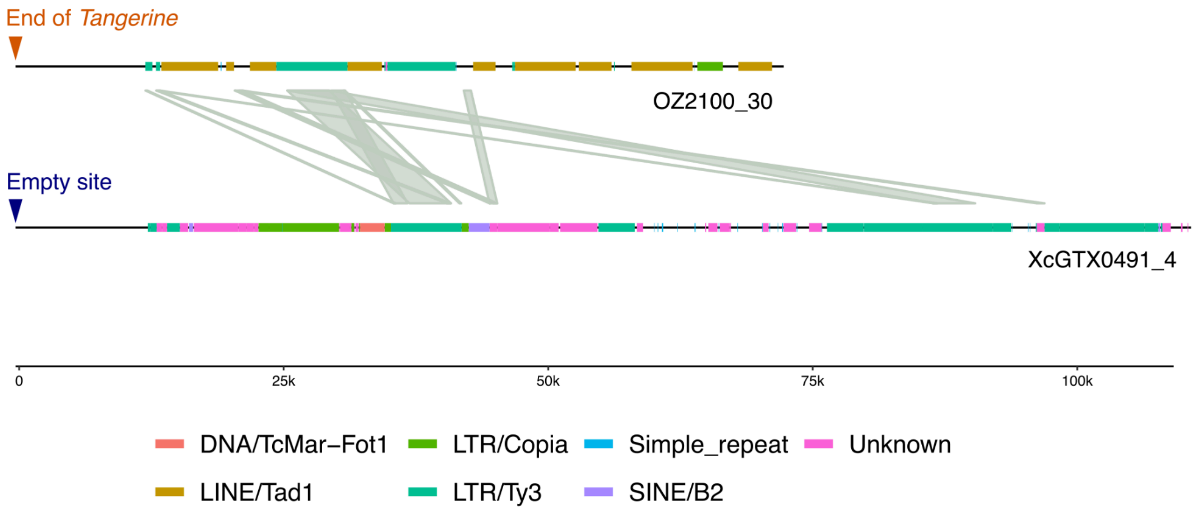


**Fig. S9. Alignment of the LRARs (large RIP-affected regions) downstream of *Tangerine* in the genome of *X. parietina* glXanPari1 (upper track) and downstream of the corresponding empty site in the genome of *X. calcicola* GTX0491 (lower track)**. Links between the fragments indicate alignments generated with BLASTn. Repeat element types are indicated with different color tracks. Repeats were annotated using Earl Grey.

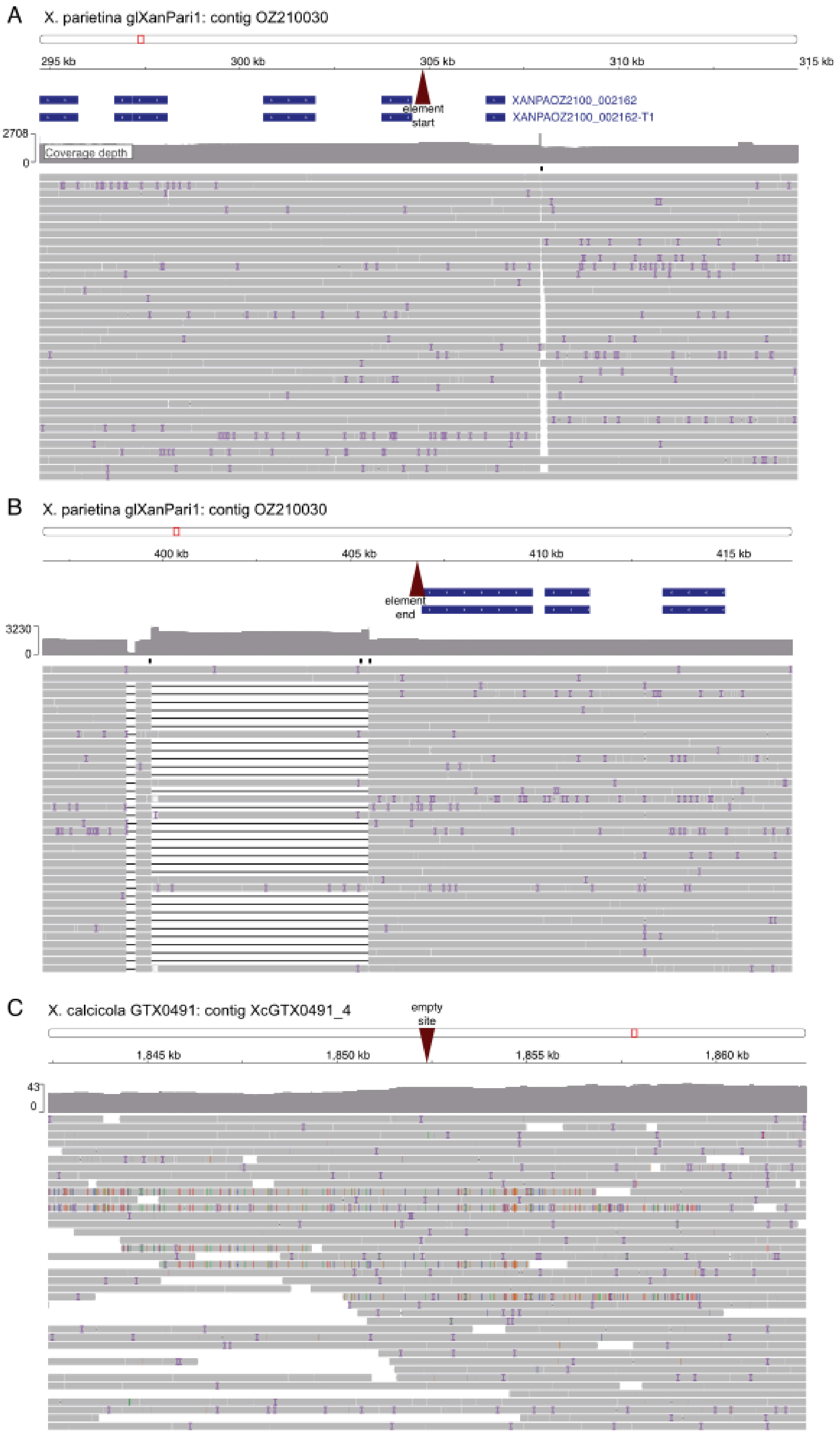
A
X. parietina glXanPari1: contig OZ210030
295 kb
300 kb
305 kb
310 kb
315 kb
element start
XANPAOZ2100_002162
XANPAOZ2100_002162-T1
2708
0
Coverage depth
B
X. parietina glXanPari1: contig OZ210030
400 kb
405 kb
410 kb
415 kb
element end
3230
0
C
X. calcicola GTX0491: contig XcGTX0491_4
empty site
1,845 kb
1,850 kb
1,855 kb
1,860 kb
43
0

**Fig. S10. Long-read data mapped to the genomic assemblies.** The upper track represents the entire contig with the red rectangle showing the position of the 20 Kbp area shown below. Gene annotations (upper track) and transcripts (lower track) are shown in blue. The grey track below shows the depth of coverage. Black rectangles under the coverage track indicate structure variants. Below, is shown a subset of reads mapped to the genome fragment. **A.** Start of *Tangerine* element on *X. parietina* glXanPari1 and 10 Kbp flanking regions. **B.** End of *Tangerine* element on *X. parietina* glXanPari1 contig and 10 Kbp flanking regions. **C.** Empty site in *X. calcicola* genome and 10 Kbp flanking regions.

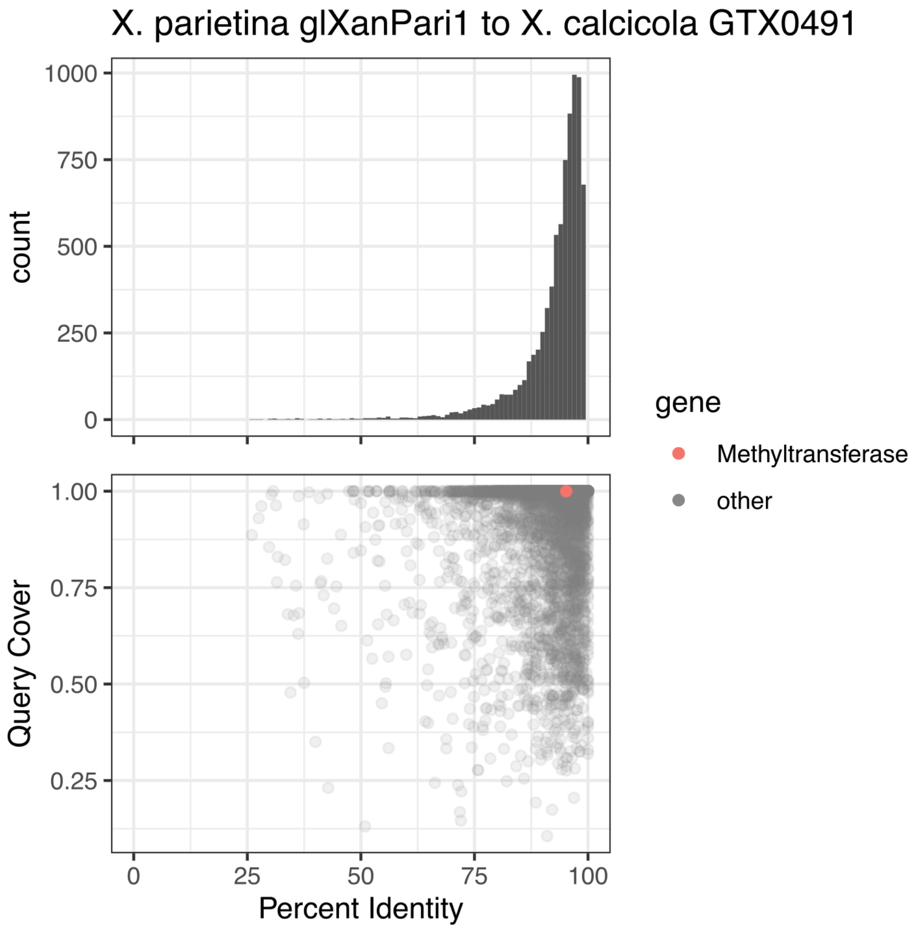


**Fig. S11. Cross-mapping of the predicted proteome of *X. parietina* glXanPari1 to *X. calcicola* GTX0491.** For each *X. parietina* glXanPari1 predicted protein, we identified the best match in the other predicted proteome. The top panel shows the distribution of percentage identity values across all matches; the bottom panel shows how the matches were positioned in relation to percentage identity and query cover. Methyltransferase is highlighted in color.

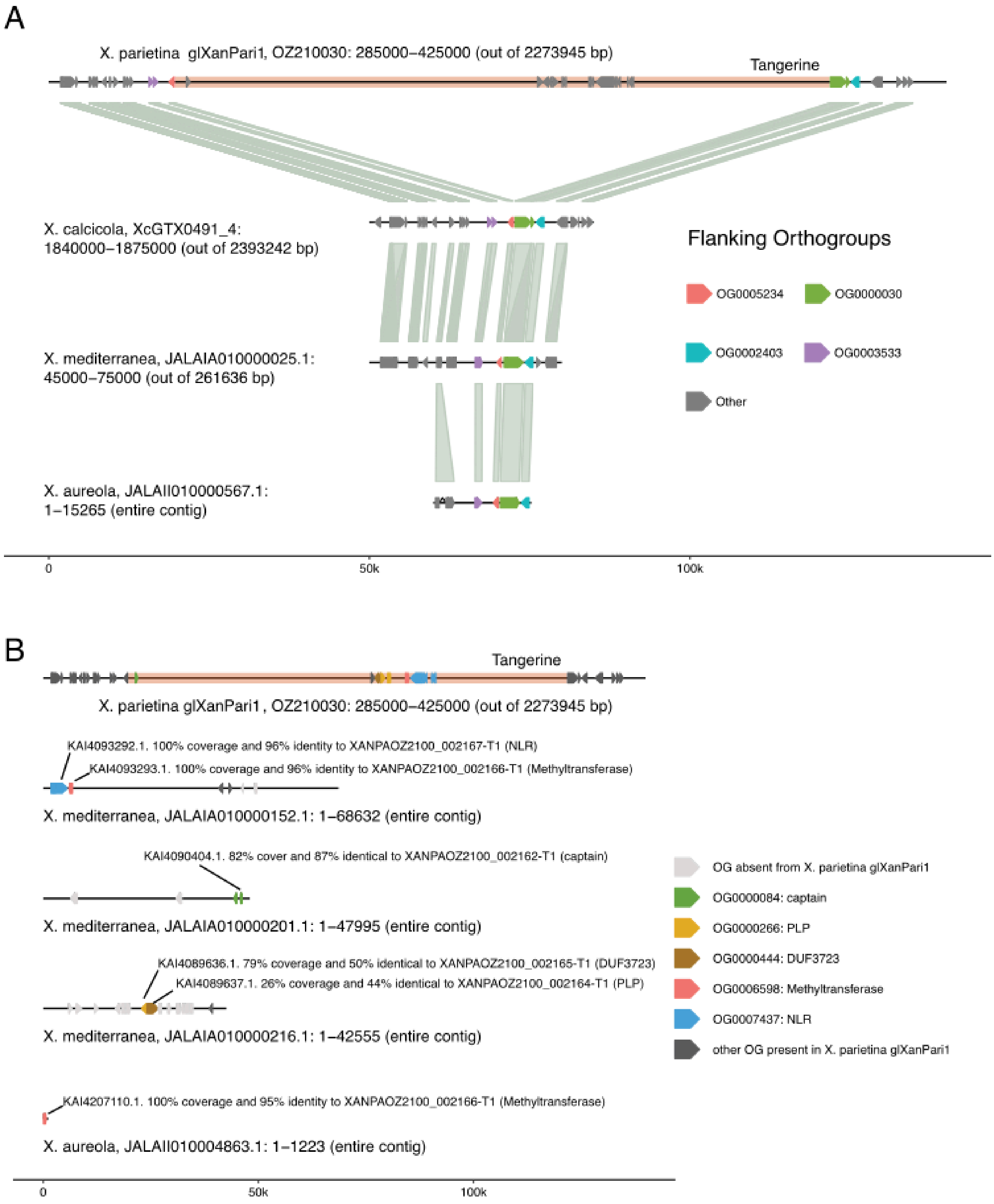


**Fig. S12. *Tangerine* in the short-read genome assemblies of *X. mediterranea* and *X. aureola*. A.** Alignment of the genomic region containing *Tangerine* in *X. parietina* glXanPari1 to the empty site in the genomes of *X. calcicola* GTX0491, *X. mediterranea*, and X. *aureola*. Genes are shown as arrows; the four flanking orthogroups are highlighted. The links connecting two genes indicate that these belong to the same orthogroup. **B.** *Tangerine*-associated genes in *X. parietina* glXanPari1 and in *X. mediterranea* and *X.*

*aureola*. Orthogroups containing the captain, accessory and cargo genes are highlighted in different colors. For each gene from *X. mediterranea* and *X. aureola* similarity scores are listed between the predicted protein and the corresponding predicted protein from *X. parietina* glXanPari1.

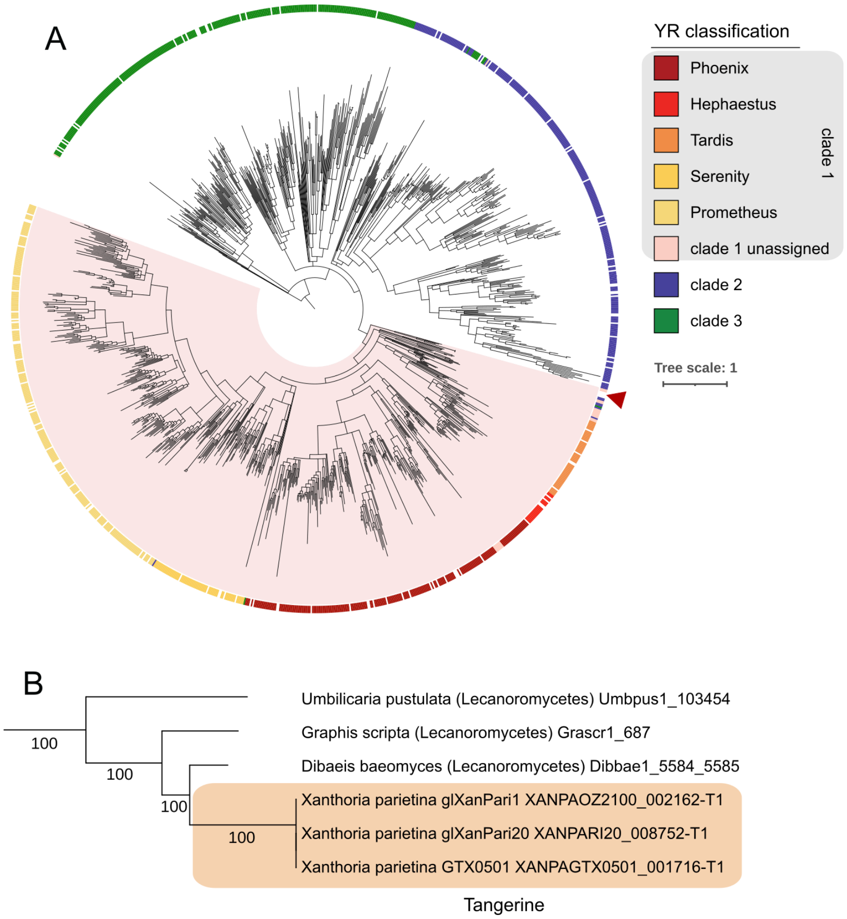


**Fig. S13. Tangerine captain in the context of the tyrosine recombinase (tyrR) typology. A.** Sequence-based captain (*Starship* tyrR) tree with the *Tangerine* captain indicated with the arrow. *Tangerine* captain sequences were aligned with 1222 predicted captains; the alignment was used to recreate maximum-likelihood phylogeny. Clades with <80 bootstrap support are collapsed. The outer color track shows captain family assignments; clade 1 is highlighted as a pink block. **B.** Magnified clade of Lecanoromycetes sequence including the *Tangerine* captain. Bootstrap support shown next to the nodes.

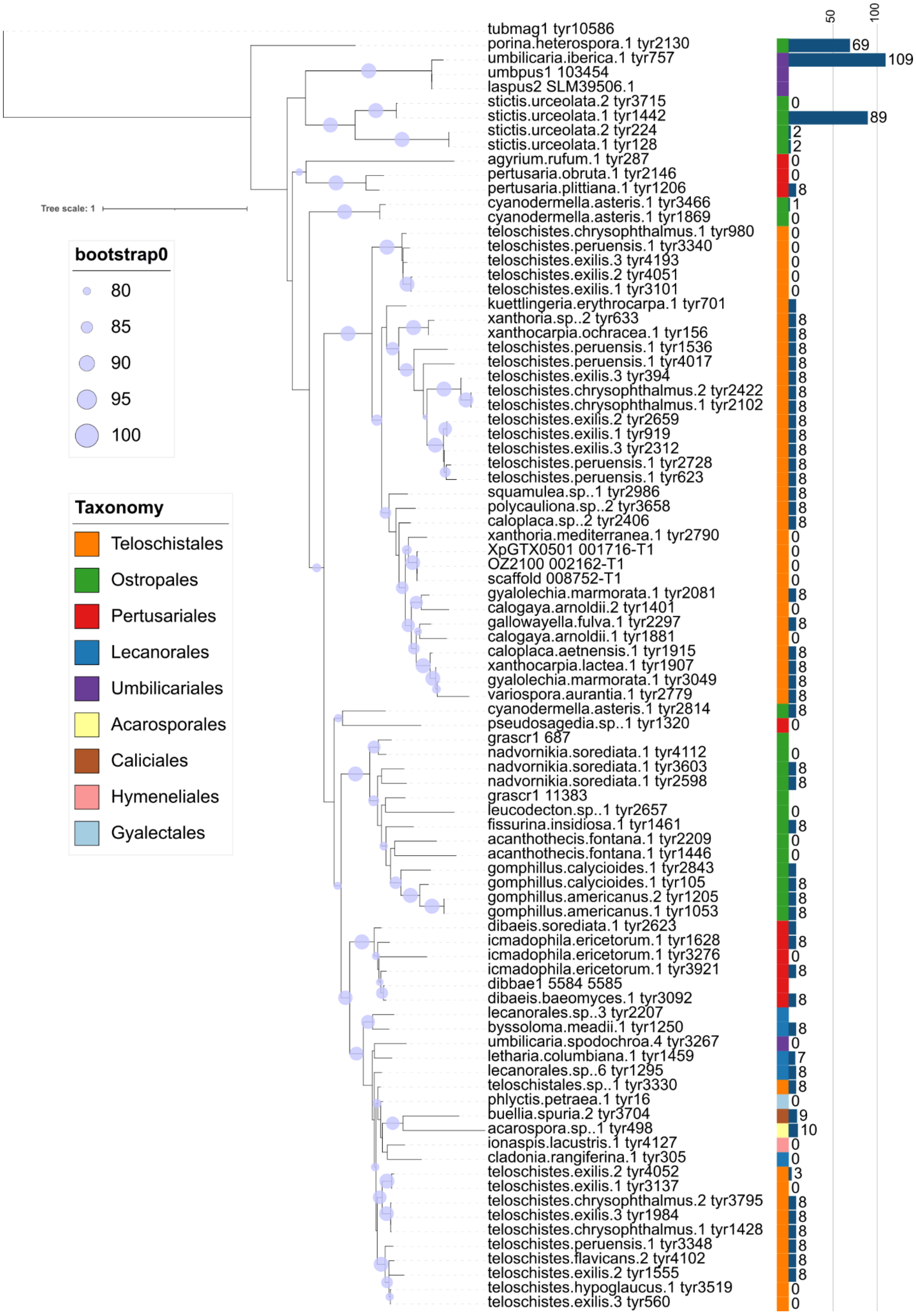


**Fig. S14. Magnified clade of the Tangerine-family showing the full set of unclustered captains, annotated based on the taxonomy of their host.** The *Tangerine* captain from *X. parietina* is indicated with the arrow. Bootstrap support shown next to the nodes. The bars to the right of the tree show the length of the CB (core-binding) domains for each captain.

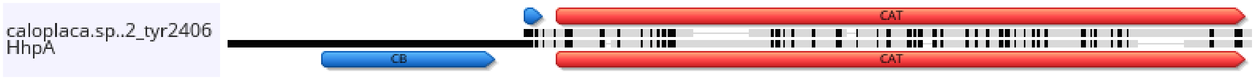


**Fig. S15. Sequence alignment of a captain from the Tangerine clade (caloplaca.sp..2 tyr2406) and Hephaestus captain HhpA.** The catalytic (CAT) domain is annotated in red. The core-binding (CB) domain of HhpA and the eight amino acid-long putative CB domain in caloplaca.sp..2 tyr2406 are annotated in blue. This putative CB domain does not align to the CB domain from HhpA. The alignment was generated using MUSCLE 5.1 and visualised in Geneious Prime 2025.2.2 (https://www.geneious.com).

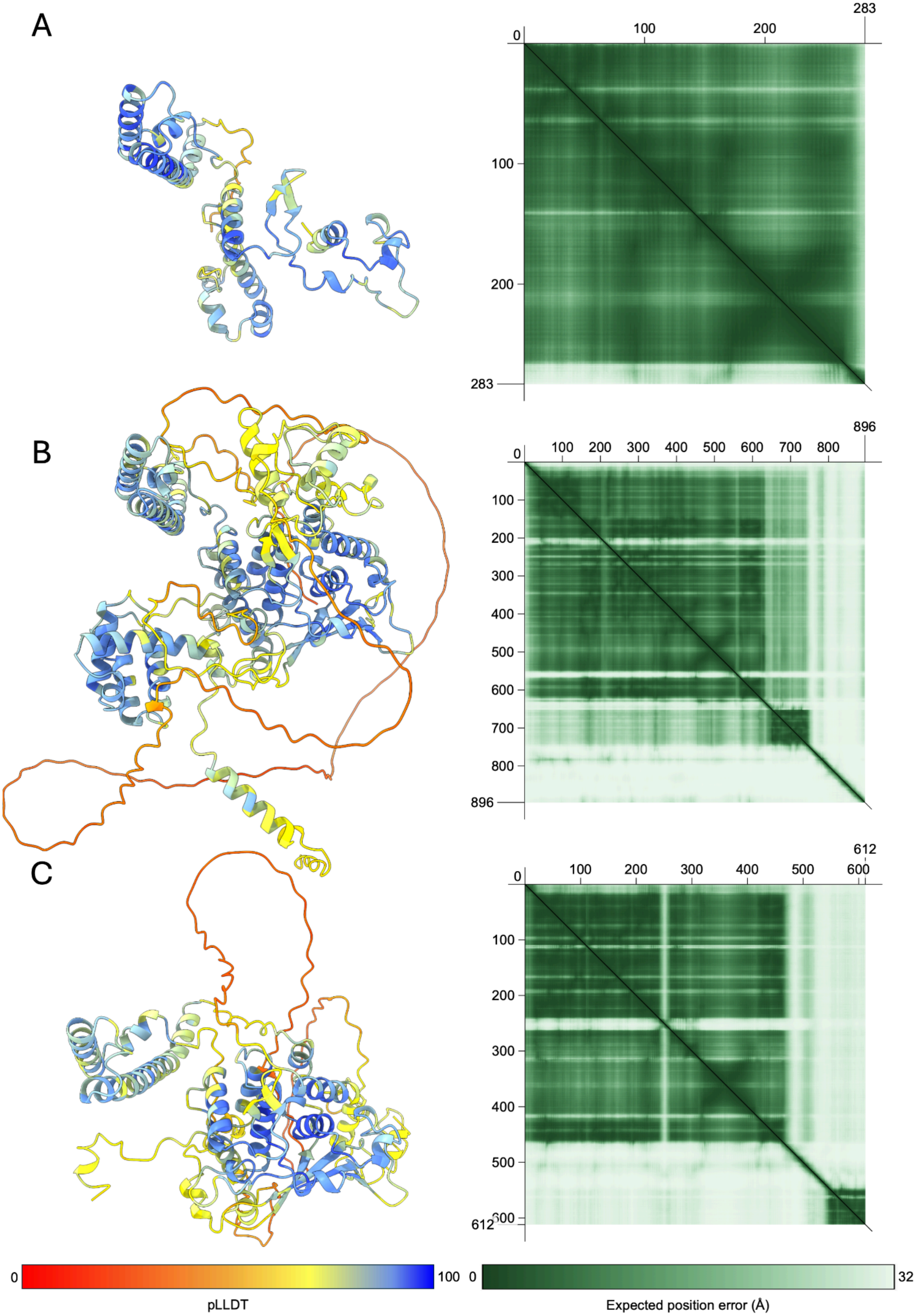
A
B
C
0
100
200
283
0
100
200
300
400
500
600
700
800
896
0
100
200
300
400
500
600
612
0
100
pLLDT
0
32
Expected position error (Å)

**Fig. S16. Prediction quality of AlphaFold3-generated monomers.** Models coloured by predicted Local Distance Difference Test (pLDDT). Associated predicted aligned error (PAE) plots, measured in Ångström (Å), included to the right of each model. **A.** *Tangerine* captain XANPAGTX0501_001716-T1 (pTM = 0.69). **B.** *Hephaestus* captain HhpA (pTM = 0.68). **C.** *Tangerine* captain caloplaca.sp..2_tyr2406 containing a section of the missing Core Binding domain (pTM = 0.68).

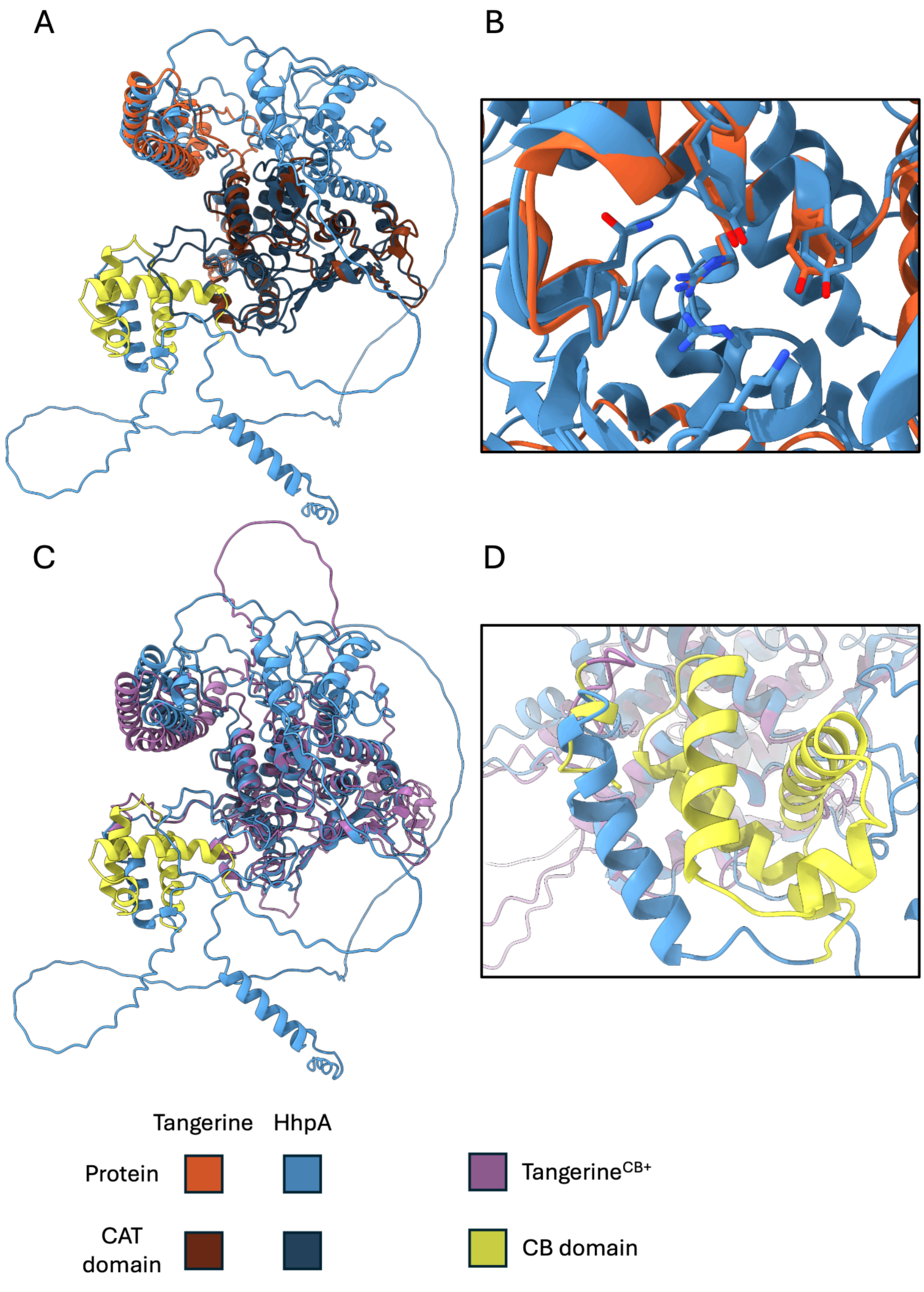

A
B
C
D
Tangerine
HhpA
Protein
CAT domain
Tangerine^CB+
CB domain

**Fig. S17. Structural superimposition of AlphaFold3-generated captains.** The catalytic (CAT) domains are differentially coloured by model, but where present the core binding (CB) domain coloured yellow. **A.** *Hephaestus* captain HhpA (blue) superimposed on top of the *Tangerine* captain XANPAGTX0501_001716-T1 (orange). Catalytic (CAT) domains highlighted in dark blue and dark orange respectively. **B.** Cut-out of the HhpA active site superimposed onto the *Tangerine* captain XANPAGTX0501_001716-T1. **C.** *Hephaestus* captain HhpA superimposed on top of the *Tangerine* captain caloplaca.sp..2_tyr2406 (purple). **D.** Cut-out of the HhpA core binding (CB) domain superimposed onto the *Tangerine* captain caloplaca.sp..2_tyr2406 with the 8 residue CB region highlighted.

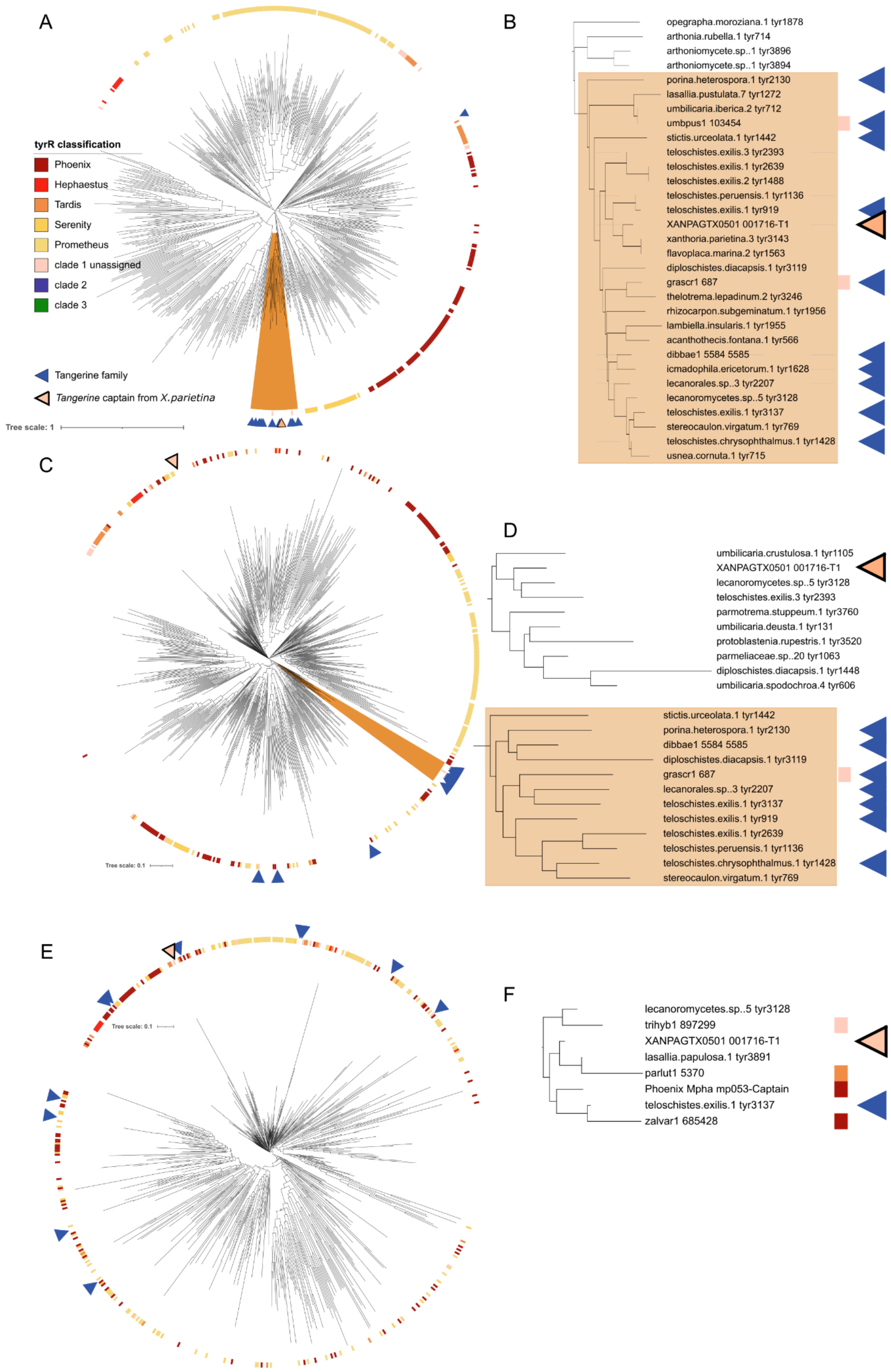
A
tyrR classification
Phoenix
Hephaestus
Tardis
Serenity
Prometheus
clade 1 unassigned
clade 2
clade 3
Tangerine family
*Tangerine* captain from *X.parietina*
Tree scale: 1
B
opegrapha.moroziana.1 tyr1878
arthonia.rubella.1 tyr714
arthoniomycete.sp..1 tyr3896
arthoniomycete.sp..1 tyr3894
porina.heterospora.1 tyr2130
lasallia.pustulata.7 tyr1272
umbilicaria.iberica.2 tyr712
umbpus1 103454
stictis.urceolata.1 tyr1442
teloschistes.exilis.3 tyr2393
teloschistes.exilis.1 tyr2639
teloschistes.exilis.2 tyr1488
teloschistes.peruensis.1 tyr1136
teloschistes.exilis.1 tyr919
XANPAGTX0501 001716-T1
xanthoria.parietina.3 tyr3143
flavoplaca.marina.2 tyr1563
diploschistes.diacapsis.1 tyr3119
grascr1 687
thelotrema.lepadinum.2 tyr3246
rhizocarpon.subgeminatum.1 tyr1956
lambiella.insularis.1 tyr1955
acanthothecis.fontana.1 tyr566
dibbae1 5584 5585
icmadophila.ericetorum.1 tyr1628
lecanorales.sp..3 tyr2207
lecanoromycetes.sp..5 tyr3128
teloschistes.exilis.1 tyr3137
stereocaulon.virgatum.1 tyr769
teloschistes.chrysophthalmus.1 tyr1428
usnea.cornuta.1 tyr715
C
Tree scale: 0.1
D
umbilicaria.crustulosa.1 tyr1105
XANPAGTX0501 001716-T1
lecanoromycetes.sp..5 tyr3128
teloschistes.exilis.3 tyr2393
parmotrema.stuppeum.1 tyr3760
umbilicaria.deusta.1 tyr131
protoblastenia.rupestris.1 tyr3520
parmeliaceae.sp..20 tyr1063
diploschistes.diacapsis.1 tyr1448
umbilicaria.spodochroa.4 tyr606
stictis.urceolata.1 tyr1442
porina.heterospora.1 tyr2130
dibbae1 5584 5585
diploschistes.diacapsis.1 tyr3119
grascr1 687
lecanorales.sp..3 tyr2207
teloschistes.exilis.1 tyr3137
teloschistes.exilis.1 tyr919
teloschistes.exilis.1 tyr2639
teloschistes.peruensis.1 tyr1136
teloschistes.chrysophthalmus.1 tyr1428
stereocaulon.virgatum.1 tyr769
E
Tree scale: 0.1
F
lecanoromycetes.sp..5 tyr3128
trihyb1 897299
XANPAGTX0501 001716-T1
lasallia.papulosa.1 tyr3891
parlut1 5370
Phoenix Mpha mp053-Captain
teloschistes.exilis.1 tyr3137
zalvar1 685428

**Fig. S18. *Tangerine* captains in the content of the structural phylogeny of tyrosine recombinases (tyrR).** We predicted a secondary structure for the *Tangerine* captain (XANPAGTX0501_001716-T1 from *X. parietina* GTX0501), manually selected representative group of *Starship* captains from clade 1 from Gluck-Thaler and Vogan (2024), and a set of *de-novo* annotated representative captains from lichen-associated fungi (Table S18) using AlphaFold, and used FoldTree to reconstruct structure-based phylogeny. The color strip represents the tyrR family according to the sequence-based classification from Vogan & Gluck-Thaler. Blue arrows show Tangerine-family captains, light-orange arrows show the *Tangerine* captain from *X. parietina*. The clades consisting primarily of the Tangerine-family captains are highlighted in orange. The tree files in the Newick format are available at the FigShare repo. **A.** Full tree based on the Fident similarity metric ('foldtree' phylogeny in the output of FoldTree). **B.** Tangerine-family clade of the tree based on the Fident similarity metric. **C.** Full tree based on the local structure alignment (LDDT). **D.** The clade containing the *Tangerine* captain from *X. parietina* (top) and the majority of the Tangerine-family clade (bottom) of the tree based on LDDT. **E.** Full tree based on the TM-scores. **F.** The clade containing the *Tangerine* captain from *X. parietina* of the tree based on the TM-scores.

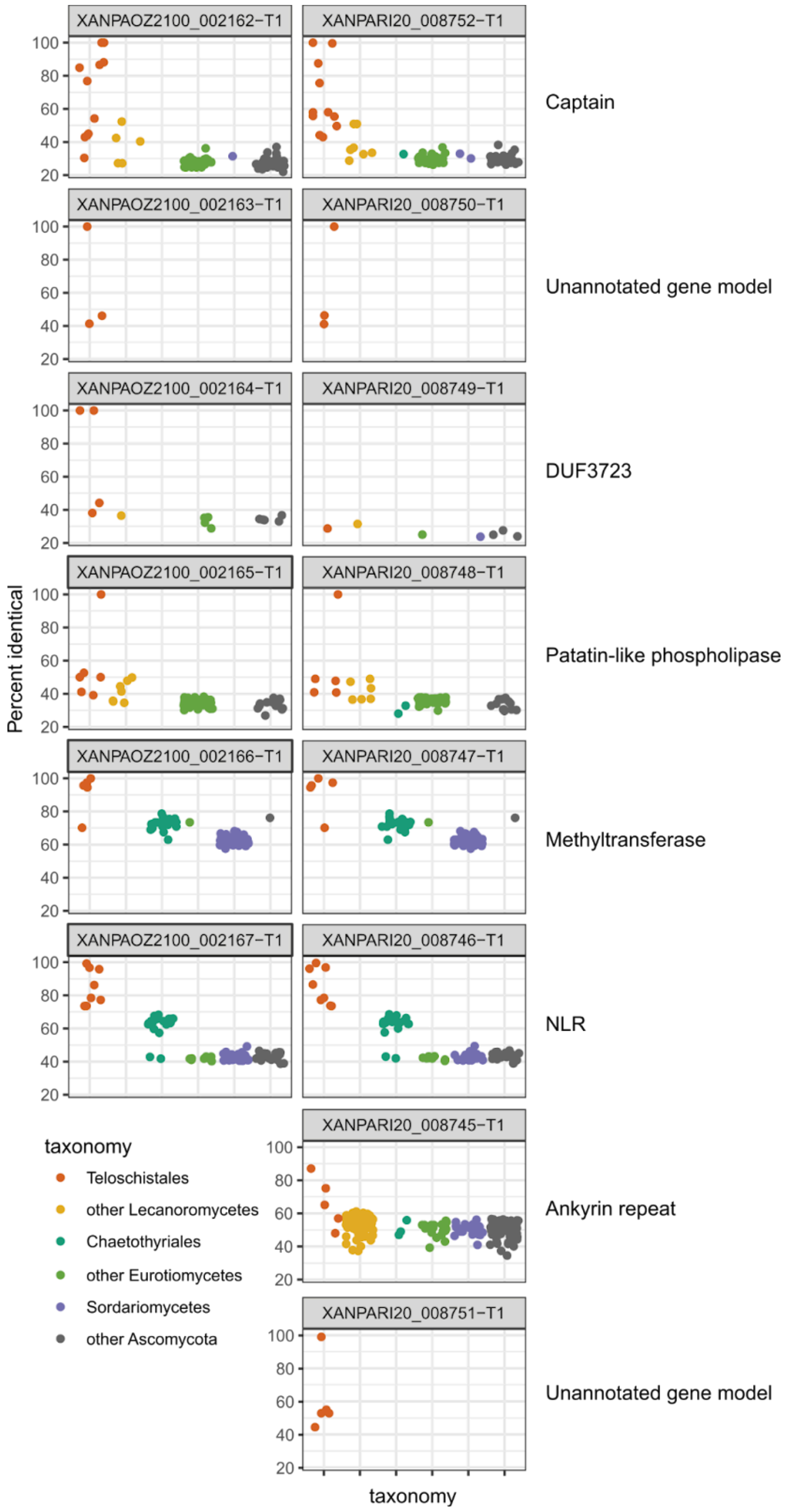


**Fig. S19. Hits to the NCBI nr database for *Tangerine*-associate genes.** We searched *Tangerine*-associate genes from *X. parietina* glXanPari1 (locus tag XANPAOZ2100) and glXanPari20 (locus tag XANPARI20) against NCBI nr, removed sequences with <50% query cover, and plotted the hits with the y-axis representing percent identity and the x-axis and color representing taxonomic origin of the hit.

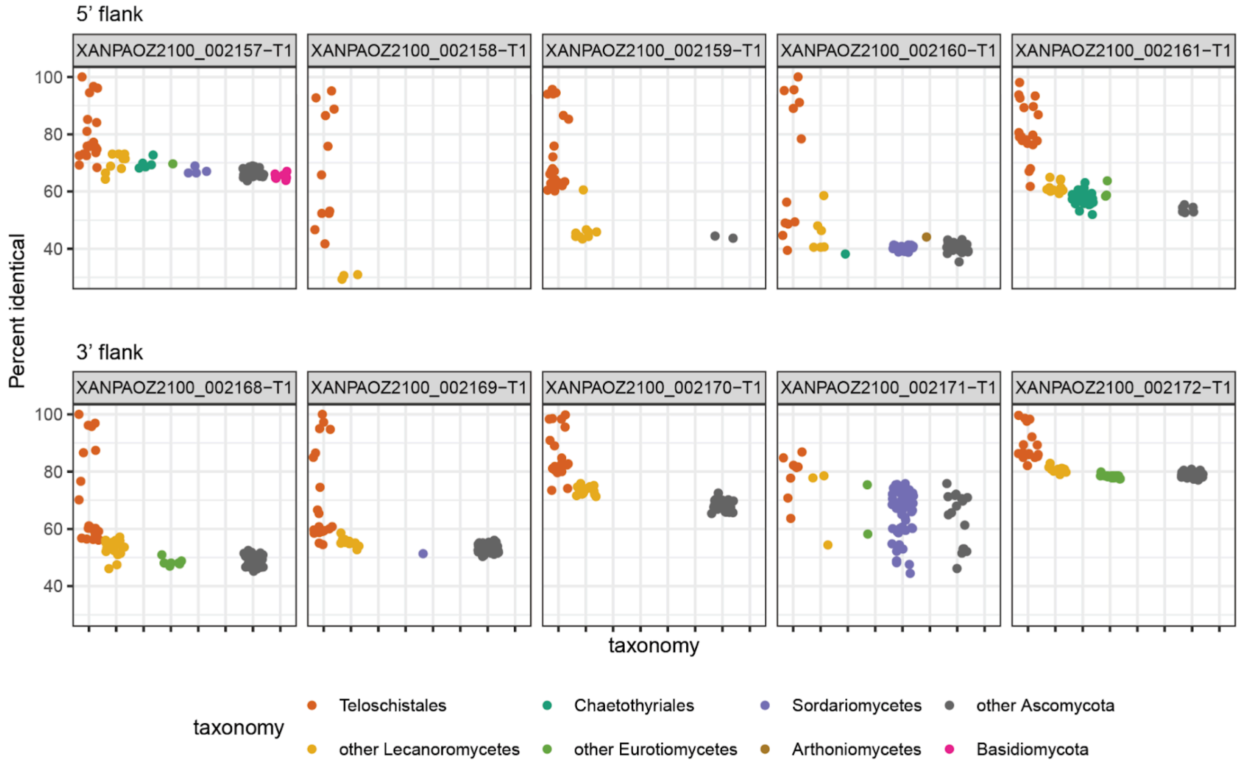


**Fig. S20. Hits to the NCBI nr database for genes in the vicinity of *Tangerine*.** We selected five gene models on the 5’ flank and five gene models on the 3’ flank of *Tangerine* in *X. parietina* glXanPari1. We searched the genes against NCBI nr, removed sequences with <50% query cover, and plotted the hits with the y-axis representing percent identity and the x-axis and color representing taxonomic origin of the hit.

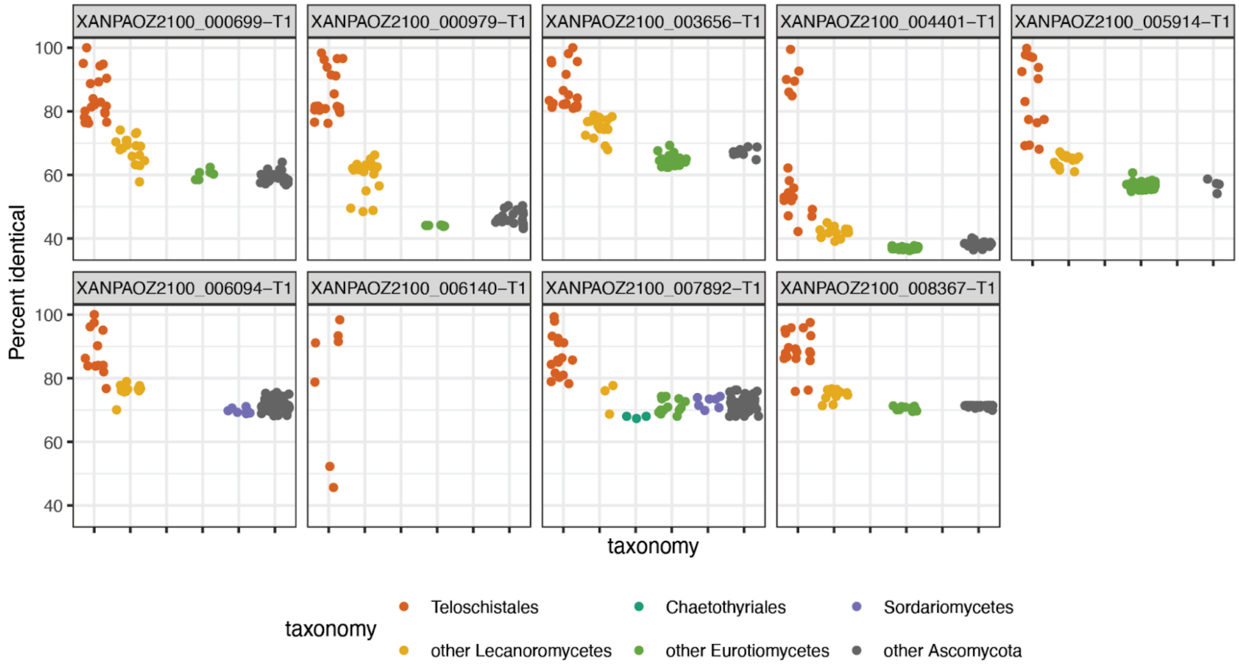


**Fig. S21. Hits to the NCBI nr database for randomly selected genes in the *X. parietina* glXanPari1 genome**. We randomly selected 10 gene models in *X. parietina* glXanPari1 and searched them against NCBI nr, removed sequences with <50% query cover, and plotted the hits with the y-axis representing percent identity and the x-axis and color representing taxonomic origin of the hit. One of the genes, XANPAOZ2100_002117-T1, did not produce any hits above the query cover threshold.

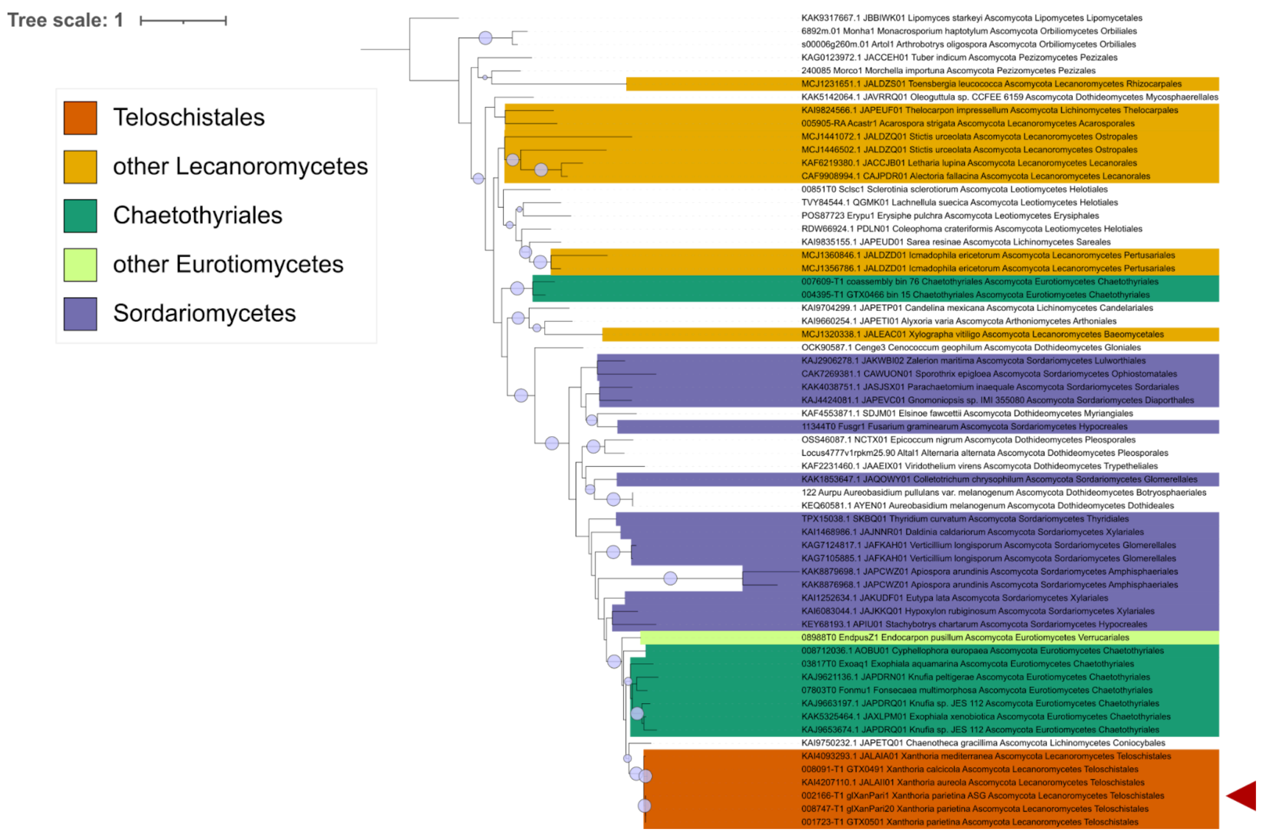


**Fig. S22. Gene tree for the orthogroup OG0006598 containing the methyltransferase.** To obtain the tree, we performed orthogroup analysis on 108 ascomycete genomes. Using the sequences for the identified orthogroup, we computed maximum-likelihood phylogeny. The tree is colored based on the taxonomic origin of sequences; bootstrap values are shown for the clades with >90 support. The sequences from *X. parietina* glXanPari1 are indicated with arrows.

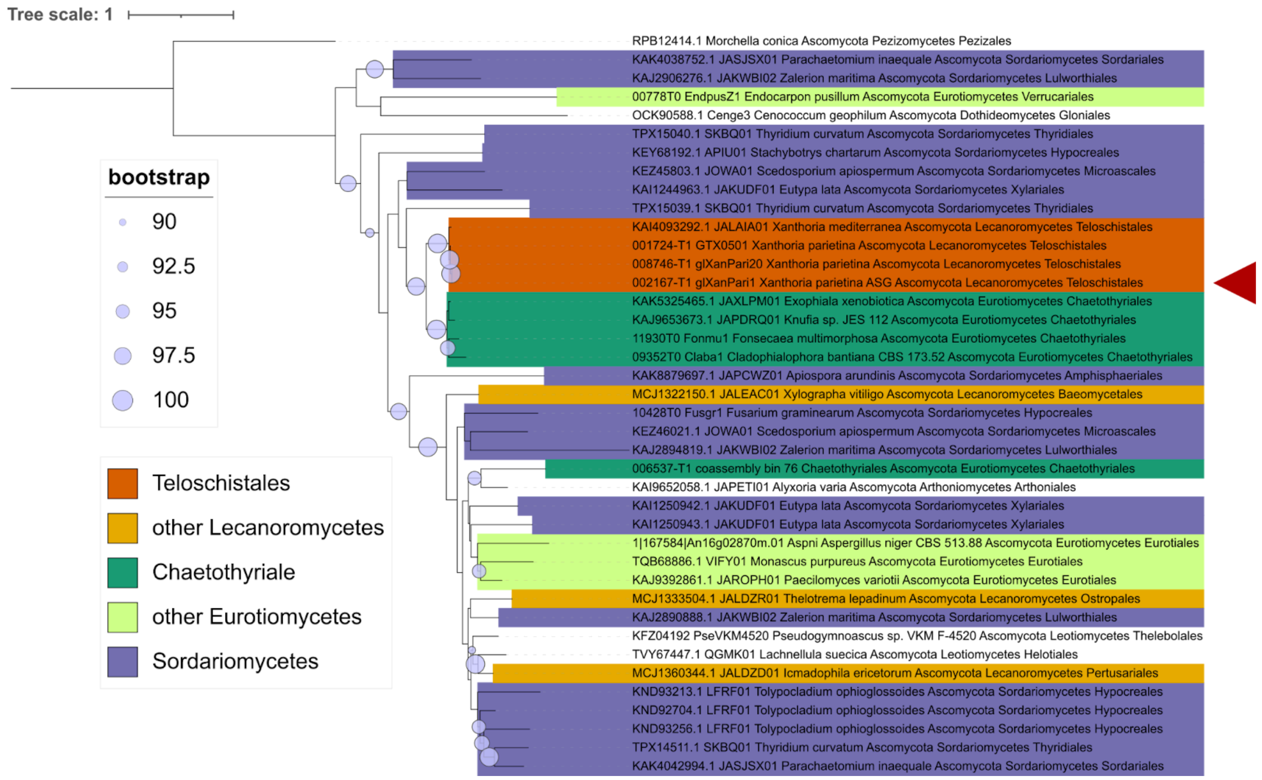


**Fig. S23. Gene tree for the orthogroup OG0007437 containing the NLR gene.** To obtain the tree, we performed orthogroup analysis on 108 ascomycete genomes. Using the sequences for the identified orthogroup, we computed maximum-likelihood phylogeny. The tree is colored based on the taxonomic origin of sequences; bootstrap values are shown for the clades with >90 support. The sequences from *X. parietina* glXanPari1 are indicated with arrows.

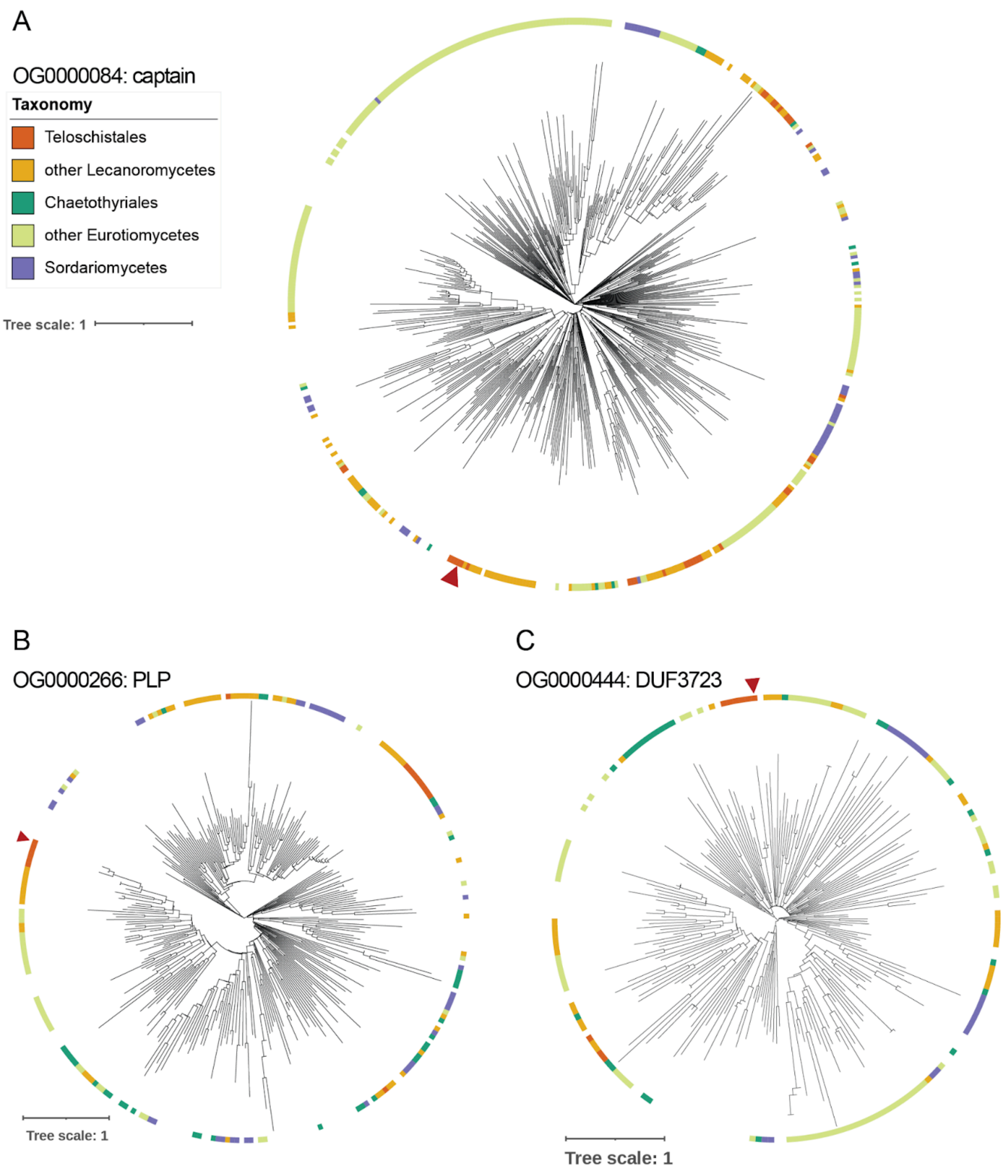


**Fig. S24. Gene trees for orthogroups containing *Tangerine*-associated genes produced by OrthoFinder. A.** Tree for OG0000084. The *Tangerine* captain (XANPAOZ2100_002162-T1) is indicated with the arrow. **B.** Tree for OG0000266. The *Tangerine* patatin-like phospholipase (PLP; XANPAOZ2100_002165-T1) is indicated with the arrow. **C.** Tree for OG0000444. The *Tangerine* DUF3723 (XANPAOZ2100_002164-T1) is indicated with the arrow. The outer ring indicates the taxonomic origin of the sequence. For DUF3723, the Teloschistales clade continuing the gene of interest is sister to a Lecanoromycete gene. For the remaining two genes, the Teloschistales clade continuing the gene of interest is nested within a larger Lecanoromycete clade.

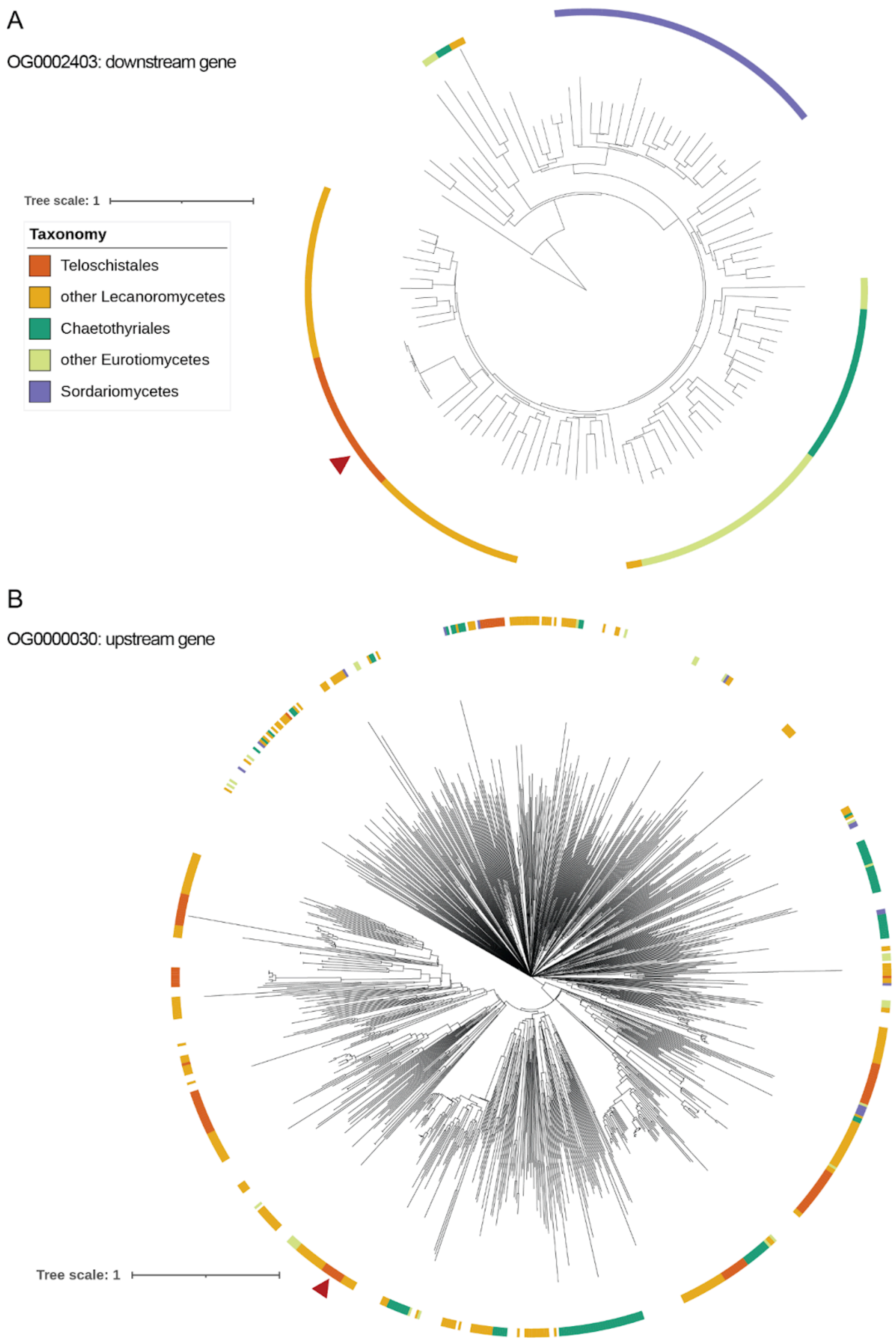


**Fig. S25. Gene trees for orthogroups containing the genes flanking *Tangerine* in *Xanthoria parietina* isolate glXanPari1. A.** Tree for OG0002403. The gene on the 3' flank of *Tangerine* (XANPAOZ2100_002168-T1) is indicated with the arrow. **B.** Tree for OG0000030. The gene on the 5' flank of *Tangerine* (XANPAOZ2100_002161-T1) is indicated with the arrow. The outer ring indicates the taxonomic origin of the sequence. For both genes, the Teloschistales clade continuing the gene of interest is nested within a larger Lecanoromycete clade.

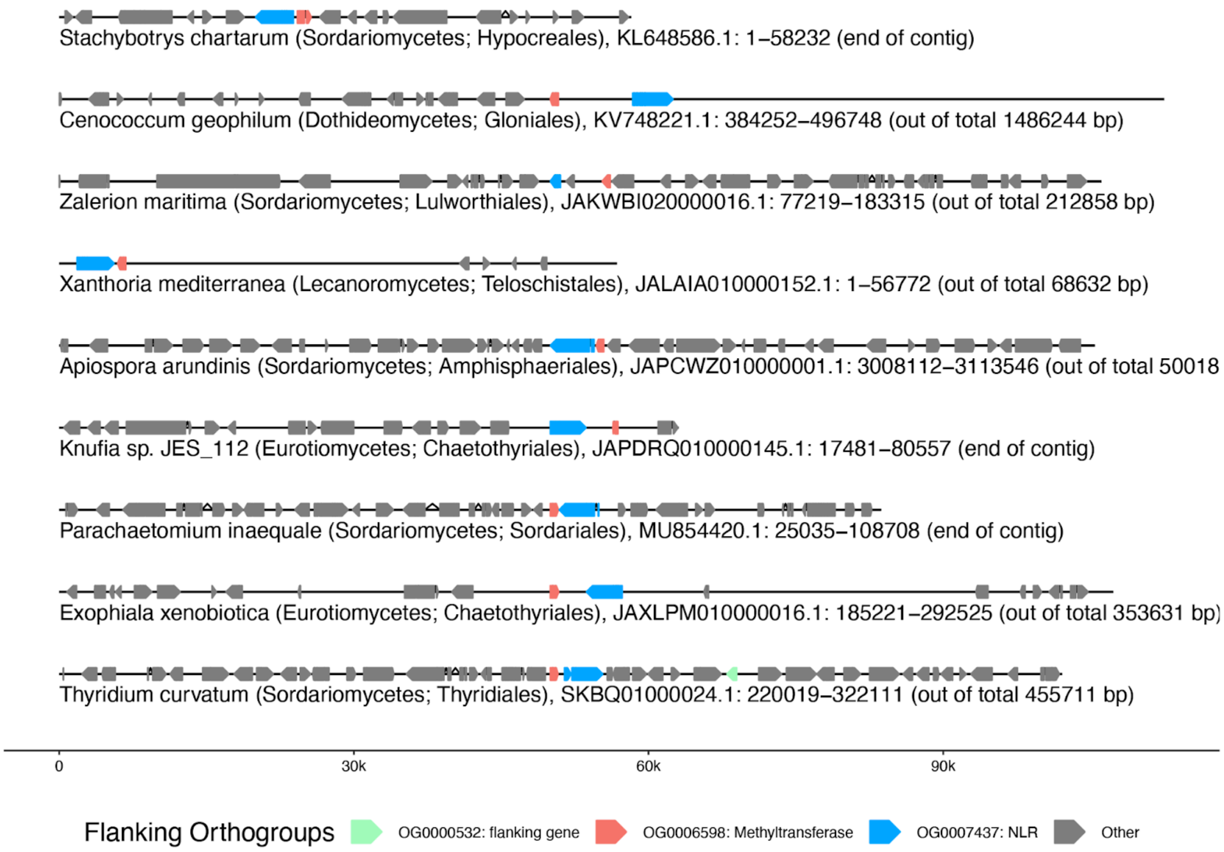


**Fig. S26. Fragments of genomes containing genes assigned to the orthogroups OG0006598 (methyltransferase) and OG0007437 (NLR) and 50 Kbp flanks**. Among the genomes included in the orthogroup analysis (Table S14), we selected genomes in which the genes assigned to the two orthogroups are placed in the same contig and separated by no more than one gene model. We plotted the contig fragments with all gene models shown as arrows. The genes assigned to orthogroups OG0006598 and OG0007437 are highlighted.

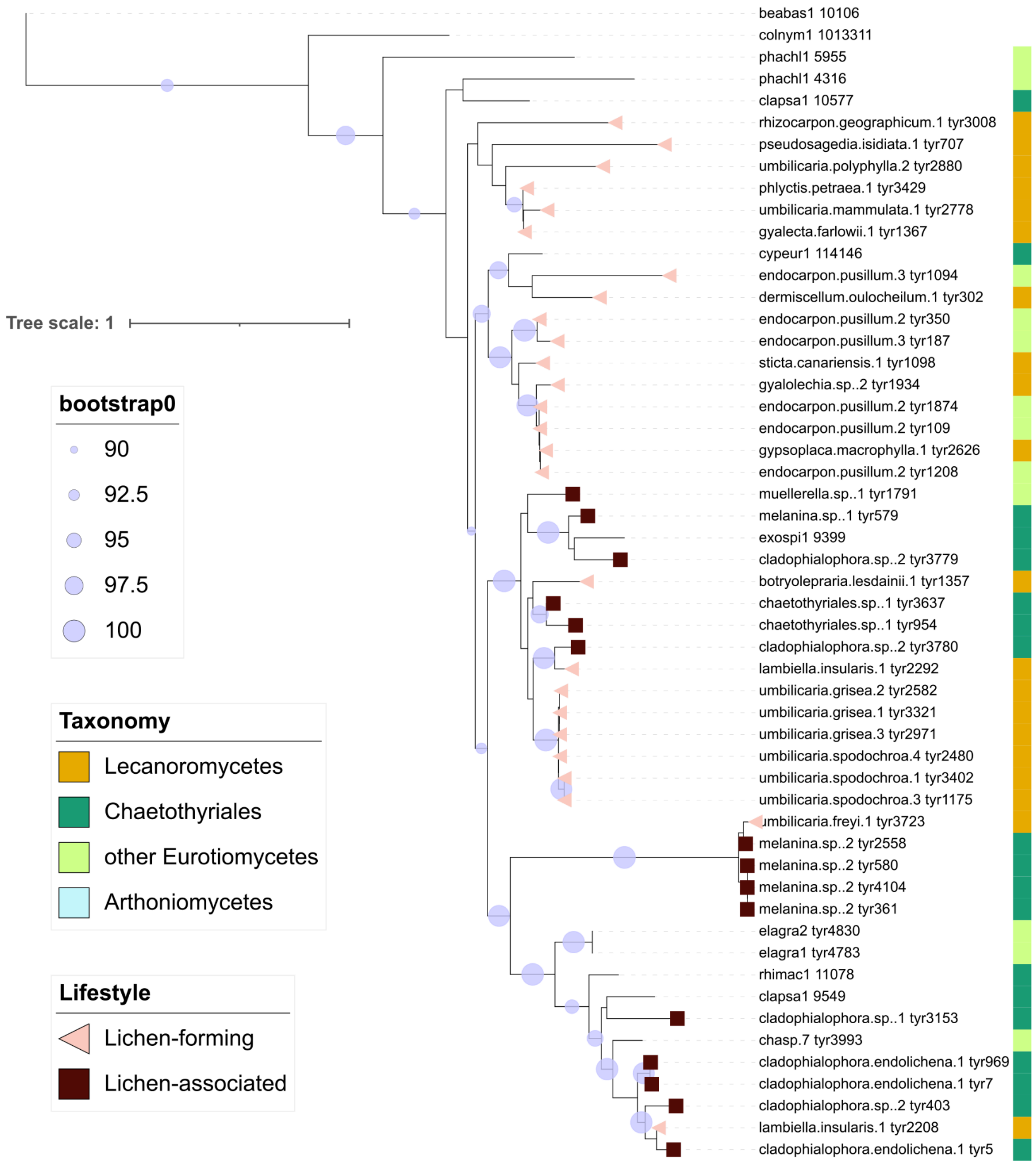


**Fig. S27. Clade A in the *Starship* tyrR phylogeny, which shows signatures of horizontal gene transfer (HGT) between distantly related lichen symbionts.** Captains within clades in the larger representative captain phylogenetic tree that showed signatures of HGT in lichen fungi were unclustered and then re-aligned; the alignment was used to recreate maximum-likelihood phylogeny. Bootstrap values are shown for the clades with >90 support. The symbols at the ends of branches indicate lifestyle: tyrRs from lichen-forming fungi (also known as mycobionts) are shown in pink triangles, tyrRs from non mycobiont lichen-associated (also known as lichenicolous or endolichenic) fungi are shown as brown squares. Sequences from non-lichen fungi lack annotation. The color strip shows the taxonomic group of the fungus. Information on the sequences is shown in Table S15.

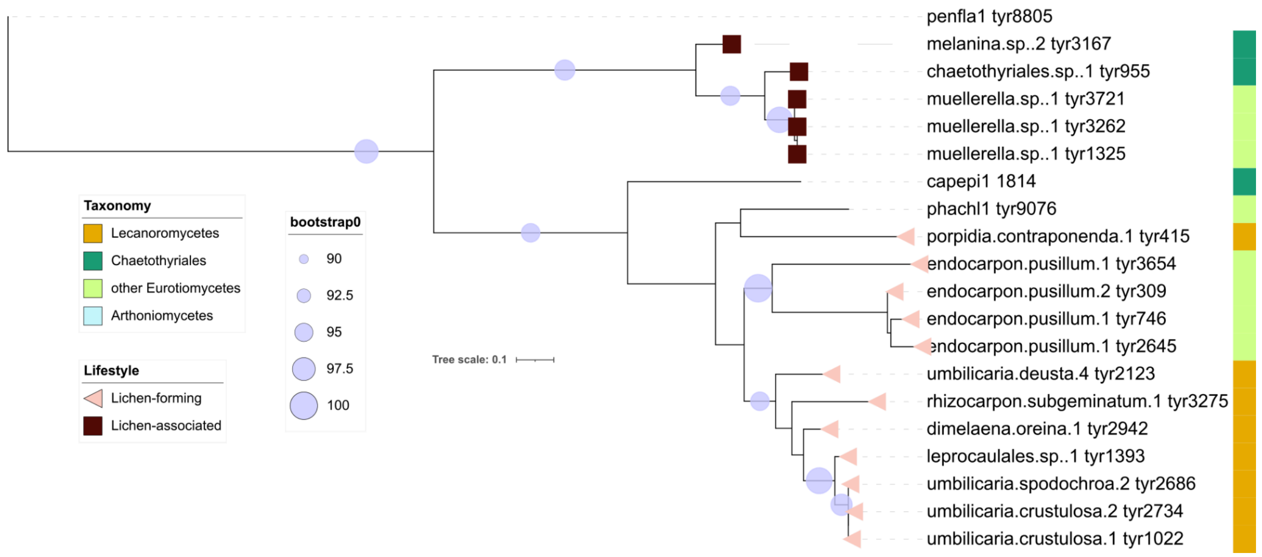


**Fig. S28. Clade B in the *Starship* tyrR phylogeny, which shows signatures of horizontal gene transfer (HGT) between distantly related lichen symbionts.** Captains within clades in the larger representative captain phylogenetic tree that showed signatures of HGT in lichen fungi were unclustered and then re-aligned; the alignment was used to recreate maximum-likelihood phylogeny. Bootstrap values are shown for the clades with >90 support. The symbols at the ends of branches indicate lifestyle: tyrRs from lichen-forming fungi (also known as mycobionts) are shown in pink triangles, tyrRs from non mycobiont lichen-associated (also known as lichenicolous or endolichenic) fungi are shown as brown squares. Sequences from non-lichen fungi lack annotation. The color strip shows the taxonomic group of the fungus. Information on the sequences is shown in Table S15.

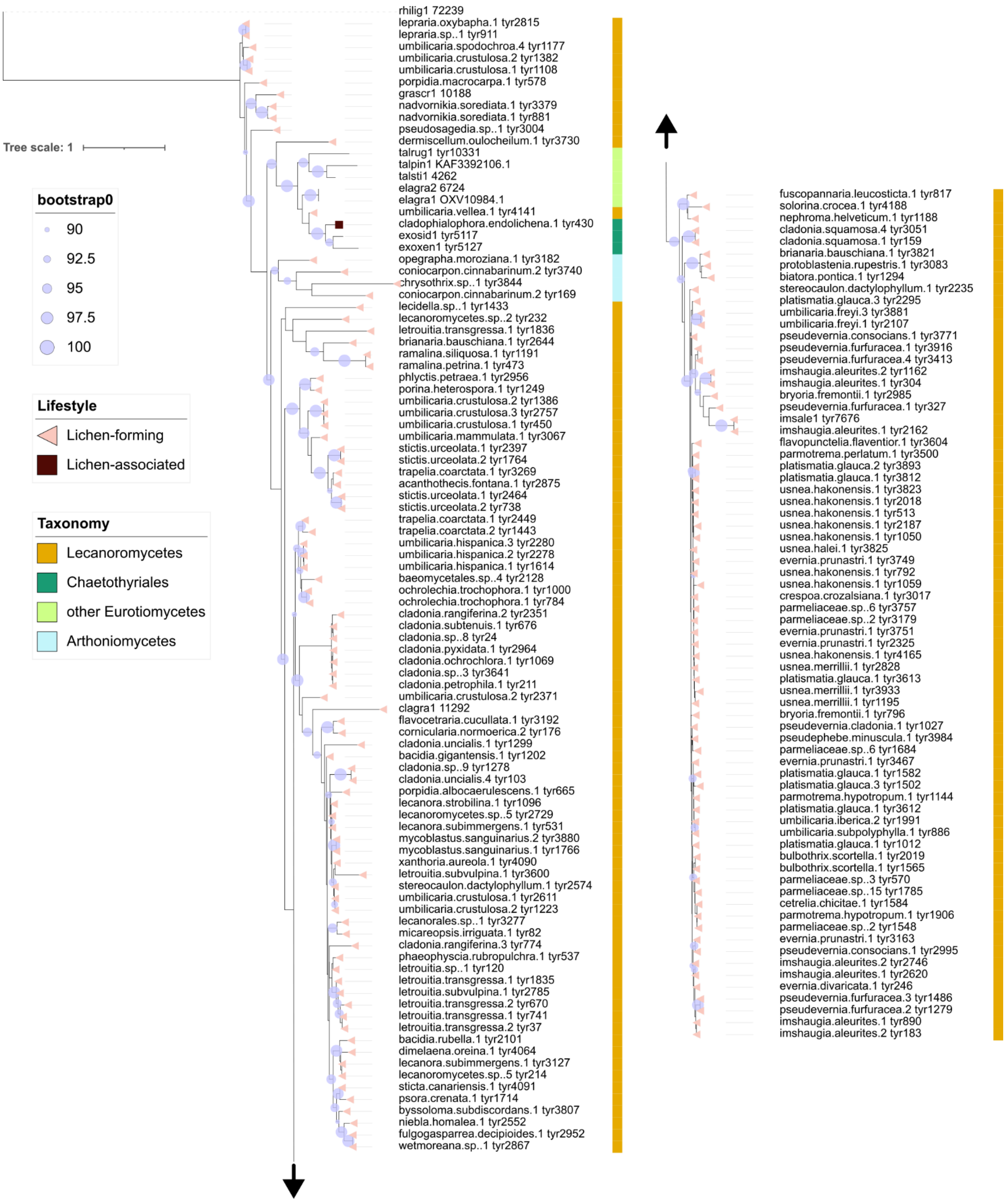


**Fig. S29. Clade C in the *Starship* tyrR phylogeny, which shows signatures of horizontal gene transfer (HGT) between distantly related lichen symbionts.** Captains within clades in the larger representative captain phylogenetic tree that showed signatures of HGT in lichen fungi were unclustered and then re-aligned; the alignment was used to recreate maximum-likelihood phylogeny. Bootstrap values are shown for the clades with >90 support. The symbols at the ends of branches indicate lifestyle: tyrRs from lichen-forming fungi (also known as mycobionts) are shown in pink triangles, tyrRs from non mycobiont lichen-associated (also known as lichenicolous or endolichenic) fungi are shown as brown squares. Sequences from non-lichen fungi lack annotation. The color strip shows the taxonomic group of the fungus. Information on the sequences is shown in Table S15.

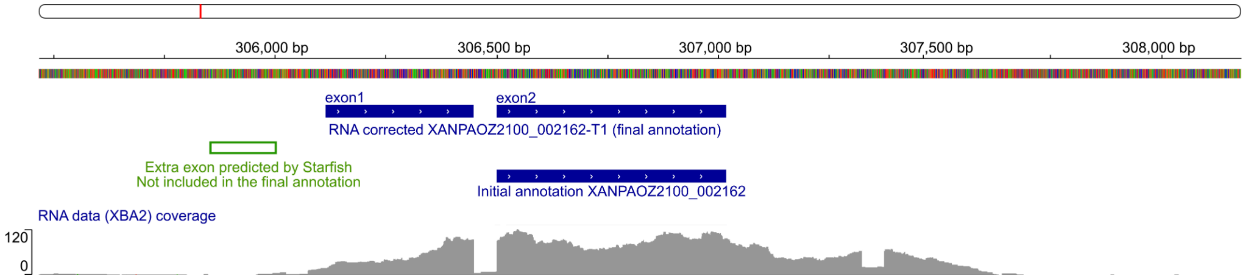


**Fig. S30. Annotation of the *Tangerine* captain XANPAOZ2100_2162 from glXanPari1 genome juxtaposed with RNA data.** Metatranscriptomic data from a *X. parietina* thallus (library XBA2) was mapped to the genome assembly using STAR. Initial annotation of the XANPAOZ2100_2162 gene model was produced by Funannotate and subsequently corrected based on the RNA mapping. When we ran the Starfish gene prediction module, a different model for this gene was produced. This model included an additional exon on the N-terminus, here shown in green. As the existence of this exon was not supported by the RNA data, it was included in the final gene model used in the downstream analyses. The data was visualized using IGV web interface.

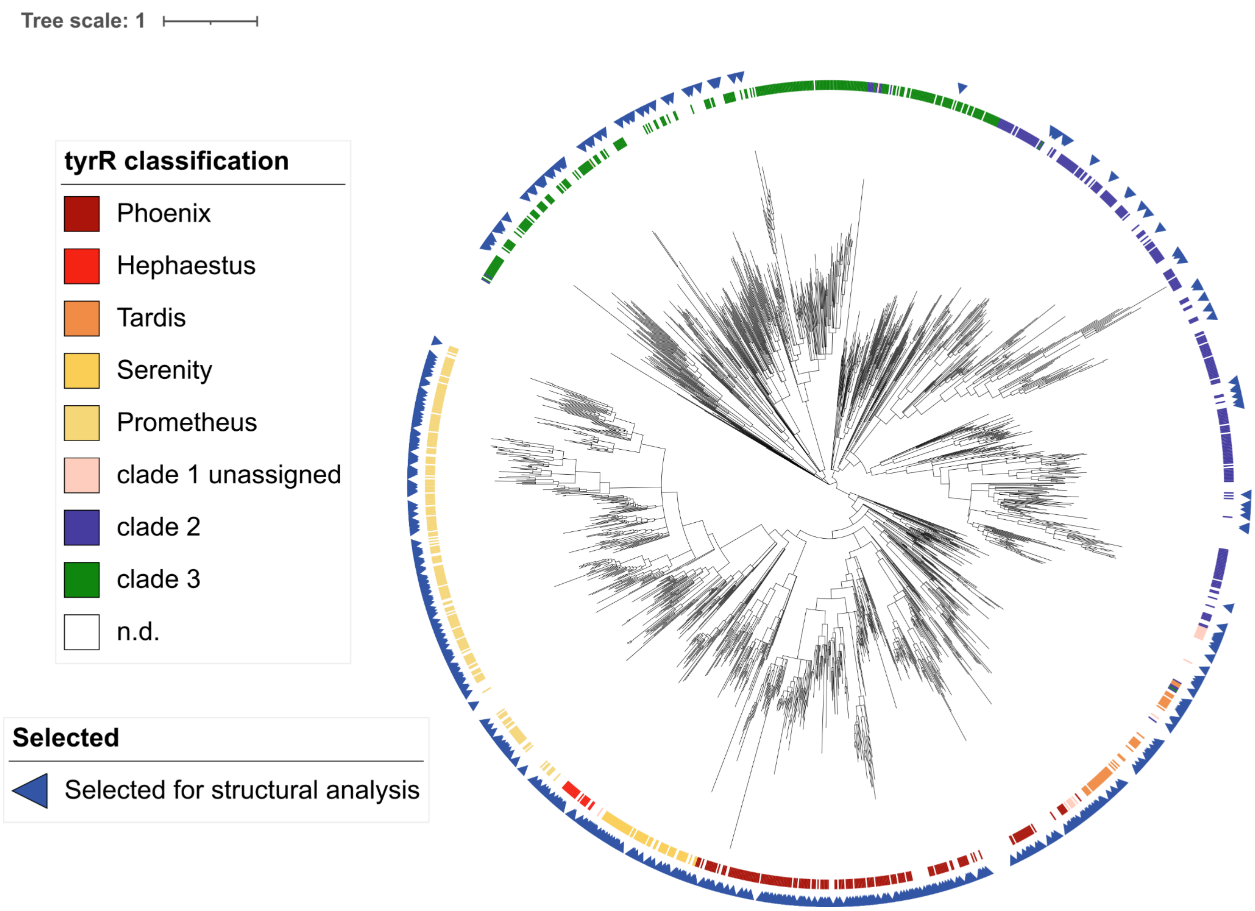


**Fig. S31. Proteins selected for structural analysis.** The sequence alignment-based tree (Fig. 3A) is shown with a color track showing family assignments for the reference tyrRs. Blue arrows show captains from lichen-associated fungi that were selected for AlphaFold structural modelling.
